# Invasive species drive polymicrobial resistance to amoxicillin in oral biofilms through β-lactamase release

**DOI:** 10.1101/2025.06.24.661089

**Authors:** Amy L. Seidel, Matthias Steglich, Taoran Qu, Amruta A. Joshi, Malina Schneider, Andrea Spaic, Ines Yang, Jasmin Grischke, Evgenii Rubalskii, Lothar Koch, Boris Chichkov, Jan Hegermann, Piotr Majewski, Elżbieta Tryniszewska, Jørgen Slots, Szymon P. Szafrański, Meike Stiesch

**Affiliations:** Department of Prosthetic Dentistry and Biomedical Materials Science, Hannover Medical School, Hannover, Germany; Lower Saxony Centre for Biomedical Engineering, Implant Research and Development (NIFE), Hannover, Germany; Department of Cardiothoracic, Transplantation and Vascular Surgery, Hannover Medical School, Hannover, Germany; Institute of Quantum Optics, Leibniz Universität Hannover, Hannover, Germany; Research Core Unit Electron Microscopy, Institute of Functional and Applied Anatomy, Hannover Medical School, Hannover, Germany; Department of Microbiological Diagnostics and Infectious Immunology, Medical University of Bialystok, Poland; Division of Periodontology, Diagnostic Sciences and Dental Hygiene, Ostrow School of Dentistry of USC, University of Southern California, Los Angeles, California, USA; Cluster of Excellence RESIST (EXC 2155), Hannover Medical School, Hannover, Germany

## Abstract

Since bacterial biofilms often cause refractory infections, antimicrobial susceptibility testing (AST) is highly desirable even for oral peri-implant biofilms. However, characterization of polymicrobial drug resistance is challenging due to high diversity and complexity of these biofilms. In this work, we developed laser-assisted AST and detected polymicrobial amoxicillin resistance in peri-implantitis. TEM-1 β-lactamase production enabled an *Enterobacter* sp. strain SPS_532 to protect its otherwise susceptible biofilm cohabitants. To understand the β-lactamase driven cross-protection in the human microbiome we aggregated genomic (n = 200,000) and patient microbial data (n = 27,000), developed a cross-protection assay, studied a representative strain collection (n = 118) and established a complex biofilm *in vitro* model (with an average of 133 species from 164 found in dental plaque). Multiple oral allochthonous species, *e.g.*, *Enterobacter*, *Klebsiella*, *Escherichia*, *Staphylococcus*, and only a single typical oral microorganism, *Haemophilus*, were able to cross-protect. Diverse *bla* genes conferred activity, via a high expression of chromosomal gene, *e.g.*, *bla*_AmpC_ gene or by the presence of plasmidic gene, *e.g.*, bla_TEM-1_ gene. Invaders not only cross-protected the biofilm from the antibiotic, but also supported expansion of opportunistic pathogens like *Fusobacterium* species. Cross-protection in complex biofilms depended on the diffusion rate and population size of the invader, which could be bio-controlled with a phage. Deciphering polymicrobial resistance might support the development of diagnostic and therapeutic approaches to combat implant-associated biofilm infections in the human mouth.

## Introduction

Due to increased demand and improved application possibilities, the number of dental implants is constantly rising, but peri-implant diseases such as peri-implant mucositis^1^ (PIM) and peri-implantitis (PI)^2^ continue to pose a serious challenge for millions of patients worldwide^3–5^. These conditions are caused by complex pathogenic bacterial biofilms and are associated with local tissue destruction, potentially leading to implant failure^6, 7^. Despite the resilience of these biofilms, clinical microbiological tests, including antimicrobial susceptibility testing (AST), are rarely used in oral peri-implant diagnostics, unlike in other areas of medicine^8–11^. Treatment of biofilm-associated diseases primarily relies on mechanical-surgical debridement and lesion drainage, often with empirically chosen broad-spectrum antimicrobials. While effective for many, this approach can fail in some patients^4^. Standardized AST, accounting for the polymicrobial nature and biofilm lifestyle of these infections, could enhance patient management by more accurately predicting *in vivo* treatment outcomes^11–14^, particularly when classical methods fall short^15, 16^. However, diversity and complexity of antimicrobial resistance profiles in oral peri-implant biofilms complicate efforts to develop tailored AST^17^. Precise, rapid, high-throughput, and programmable bioprinting of biofilm fragments onto diverse culture media presents a promising avenue for advancing AST application^18^.

β-lactam antibiotics, such as amoxicillin, have been widely used to combat biofilm-associated oral infections. But ever since their discovery, bacterial resistance has posed difficulties during treatment. β-lactamases, enzymes that hydrolyze and inactivate β-lactams, are major drivers of resistance^19^ and can be found in saliva^20^ and in biofilms from polymicrobial infections which respond poorly to β-lactam treatment, *e.g.*, periodontitis, sinusitis, and others^21–23^. Metagenomic studies have frequently detected β-lactamase genes in the oral cavity and have been able to associate their occurrence with specific species^24–29^. A metatranscriptomic study linked β-lactamase expression to periodontal disease, while isolation experiments confirmed the presence of β-lactamase-producing bacteria in 50 – 70% of patients with general, severe or refractory periodontitis^21, 30–33^. Resistance to amoxicillin has been frequently demonstrated in typical oral opportunistic pathogens^31, 34–36^ and invasive species, with β-lactamases being detectable in a large number of isolates^21, 37–41^. These enzymes exhibit remarkable diversity in substrate specificity, inhibitor sensitivity, expression levels, co-occurrence, transferability, and taxonomic distribution. While the genetics and regulation of β-lactamases have been well studied in single species infections, their role in complex biofilm-associated infections is poorly addressed^13, 42^. Interspecies cross-protection based on β-lactamase production by oral isolates has been reproduced in a mice abscess models^43^. Notably, potent allochthonous β-lactamase producers have been identified in oral pathologies, including refractory cases^21, 44, 45^. Targeting invasive opportunistic pathogens while preserving the benign oral flora is challenging, as many of these taxa exhibit high resistance to antimicrobials and antiseptics^46, 47^. Additionally, their ability to withstand harsh conditions enables them to persist in the patient’s environment, posing a risk of re- or cross-infection^48–50^. Bacteriophages and their enzymes have been proposed as potential adjuncts to antimicrobial and antiseptic treatments^33, 34^. However, the exact mechanisms underlying the cross-protection process and its clinical significance as well as the pathogenicity, virulence and ecological role of the producers in these infections remain largely unexplored ^51–53^ Additionally, the efficacy of phages against complex oral biofilms is still uncertain.

To address these challenges, we develop a laser-assisted antimicrobial susceptibility testing (AST) method and identify polymicrobial amoxicillin resistance in peri-implantitis, driven by TEM-1 β-lactamase production in *Enterobacter* sp. To further investigate β-lactamase-driven cross-protection within the human microbiome, we analyze genomic (n = 200,000) and microbial (n = 27,000) datasets. We execute cross-protection assays, involving over 100 representative strains or clinical samples, along with complex biofilm model experiments, to reveal key producers and mechanisms underlying this phenomenon and to investigate the effectiveness of bacteriophages. Our goal is to integrate the advantages of bacteriophages further into the medical environment and pave the way for innovative therapies in times of antibiotic resistance. This study propels efforts to gain deeper understanding of polymicrobial resistance and could enhance diagnostics and treatment strategies for implant-associated biofilm infections.

## Results

### Laser-assisted bioprinting of clinical biofilms facilitated detection of polymicrobial amoxicillin resistance conferred by *Enterobacter* sp. producing a TEM-1 β-lactamase

We adapted laser-assisted bioprinting to study antibiotic resistance in complex peri-implant biofilms (**Fig. 1**). This method allows precise deposition of picoliter-sized biofilm fragments into organized colony arrays (**Fig. 1a**), where local interactions occur through metabolite and enzyme diffusion. Submucosal biofilm samples from refractory peri-implantitis were printed onto media containing common dental antibiotics, either incorporated into the medium or applied via antimicrobial susceptibility test disks to create a concentration gradient.

**Fig. 1.**
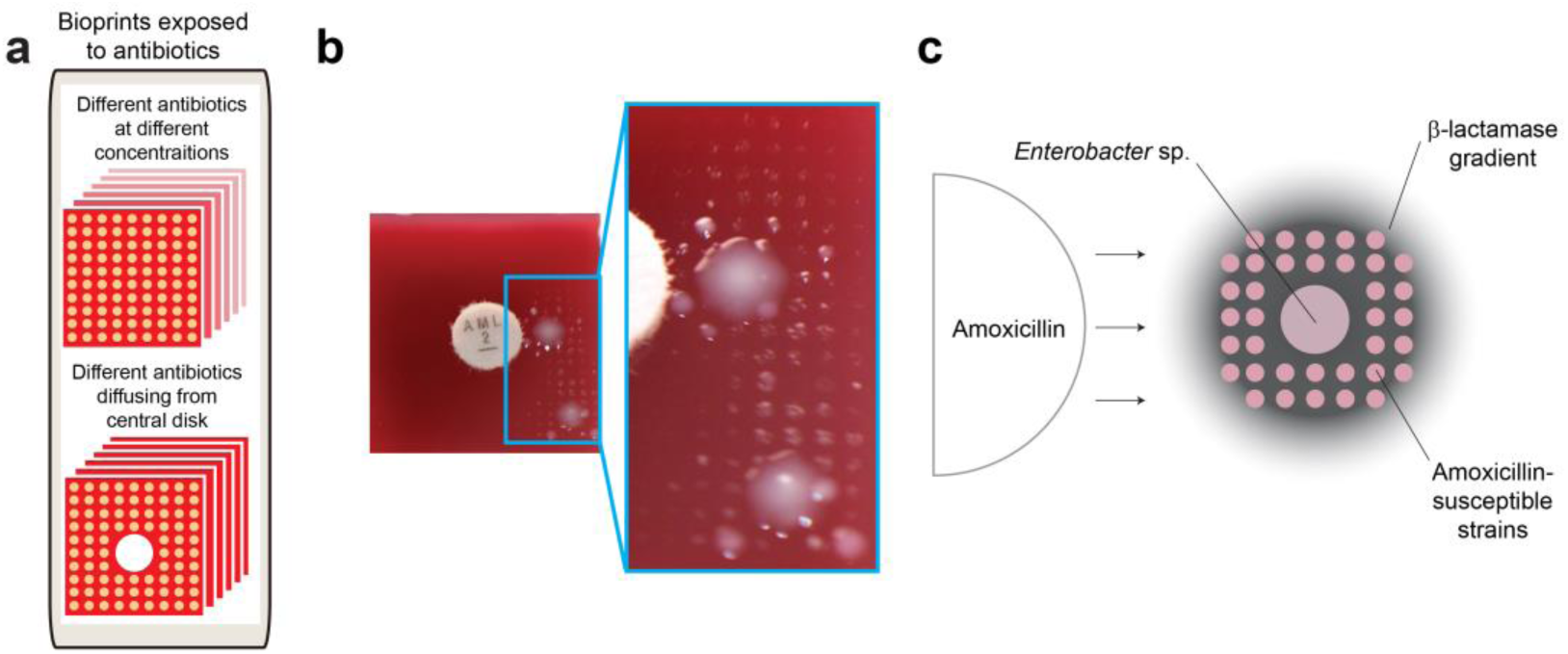
Laser-assisted bioprinting of clinical biofilms detected polymicrobial resistance driven by a β-lactamase **a** Method for laser-assisted antimicrobial susceptibility testing of oral biofilms. **b** Detection of polymicrobial resistance to Amoxicillin in a case of severe chronic peri-implantitis. **c** Mechanism of detection of polymicrobial resistance to Amoxicillin involving a β-lactamase release.

On both control media and media with a metronidazole disk biofilm arrays showed unaffected growth of around 400 of the technically possible 440 colony biofilms (**Fig. S1**). Inhibition zones appeared around tetracycline and ciprofloxacin disks, while amoxicillin disks typically inhibited growth. Occasionally, a satellite growth pattern was observed next to amoxicillin disks (**Fig. 1b, Fig. S1**), indicating polymicrobial resistance mediated by a diffusible agent. Polymicrobial resistance patterns consisted of a central colony formed by *Enterobacter* sp., surrounded by colonies of taxonomically diverse susceptible strains. Cross-protection interactions were reproduced using pairs of isolates (*e.g.*, a helper *Enterobacter* sp. SPS_532 strain and a co-isolated susceptible *Lactobacillus* sp. SPS_449 strain; **Fig. S2a**). The diffusible agent responsible for the cross-protection activity was heat- and protease-sensitive (**Fig. S2b** and **S2c**), and was identified as a TEM β-lactamase without extended-spectrum activity (non-ESBL) using chromogenic substrates, antibiotic testing, β-lactamase inhibitors and stimulators, as well as PCRs (**Fig. S2d** – **S2g**). The helper strain did not hyper-produce AmpC β-lactamase (**Fig. S2f** and **S2g**). However, colonies of hyper-producers were obtained when the helper strain was exposed to cefpodoxim in combination with an AmpC stimulator in absence of AmpC inhibitors (**Fig. S2g**). In summary, *Enterobacter* sp. (*bla*_AmpC_^wt^, *bla*_TEM-1_) conferred cross-protection to diverse biofilm co-inhabitants in peri-implantitis through the release of TEM-1 β-lactamase (**Fig. 1c**).

### Gammaproteobacteria and Bacteroidia are the major β-lactamase producers in human microbiome

Discovery of β-lactamase-driven cross-protection in refractory biofilm infections raised questions about β-lactamases producing human-associated microorganisms. Analyses of over 150,000 genomes from metagenomes covering different age groups, geography, and lifestyles^54^ revealed that β-lactamase producers were present in 48% of approximately 9.5 thousand human samples and in 5.6% of metagenomic assemblies. It should be noted that these samples mostly represented the gut flora of healthy adults, from few countries, with westernized lifestyle (**Fig. S3a** – **S3e**). In total, 8,604 β-lactamase-related genes were detected in 290 species bins (**Fig. 2a**). Genomes of *Escherichia* from the γ-Proteobacteria class and *Bacteroides* from the Bacteroidia class encoded 83% of those features. Other members of the aforementioned classes (*e.g.*, *Prevotella*, *Klebsiella* and *Enterobacter*) and other classes (*e.g.*, *Capnocytophaga* from Flavobacteriia) were also identified as prominent producers. Class A β-lactamase (*e.g.*, CblA, Cfx), Penicillin-binding proteins (PBPs) and class C β-lactamase (AmpC-related) were the most common gene products (**Fig. 2b**).

**Fig. 2.**
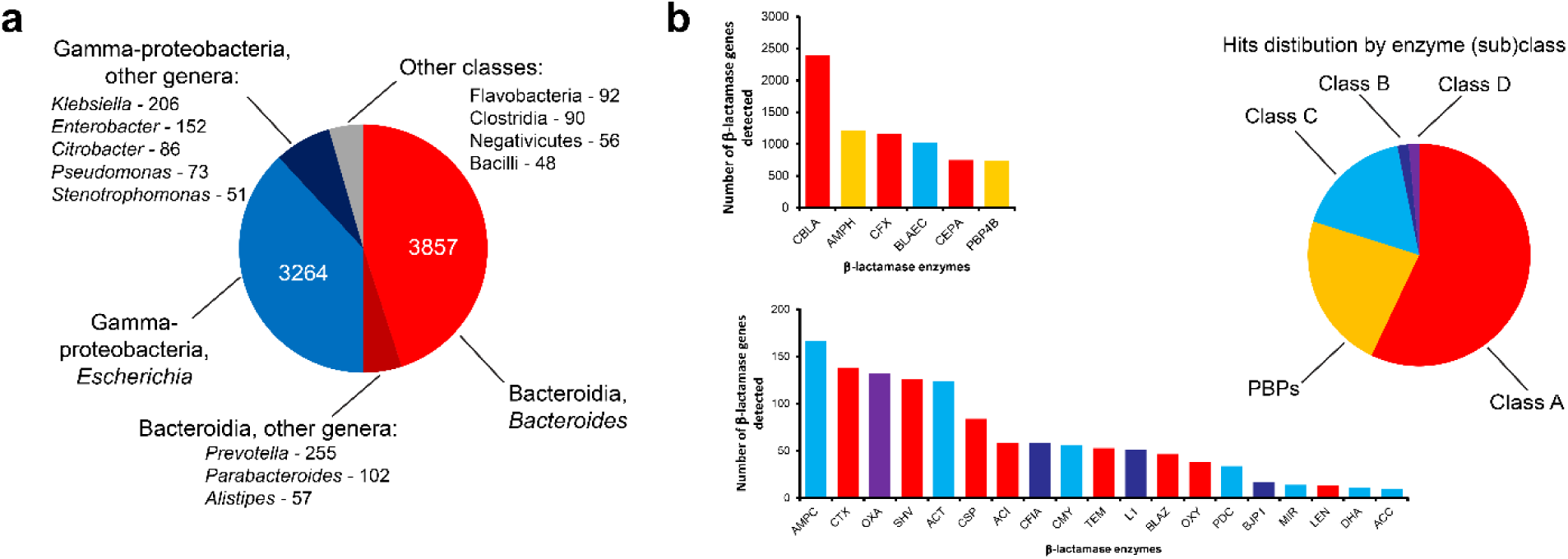
Major β-lactamase producers and their enzymes in human microbiome. **a** Major β-lactamase producers in human microbiome samples. **b** Major β-lactamases detected in human-associated metagenomes.

### In the oral cavity, allochthonous species and only few commensals produce β-lactamases

In the published metagenomes representing 783 samples from the human oral cavity (mostly collected from supragingival plaque or tongue dorsum of healthy adults), we detected only 164 β-lactamase-related genes representing 9% of all resistance marker genes. While *Prevotella* species were the major producers, we detected diverse minor producers as well (**Fig. S3f**). β-lactamase-related genes were usually bla*_CfxA_*-related and occasionally represented *bla_ampC_*, *blaZ*, bla*_OXA_*, bla*_SRT_* or *bla_TEM_*.

In order to further characterize the microbial populations of β-lactamase producers in human oral cavity, we supplemented the metagenomics results with data for representative genomes (n = 1,527) from the expanded Human Oral Microbiome Databases (www.homd.org). 286 β-lactamase-related genes were detected in 151 genomes (*i.e.*, in 10%). 32.5% hits were detected in all five major reference databases (*i.e.*, argannot, megares, card, ncbi and resfinder), 38.5% hits were detected independently using two to four databases and the remaining 29% came from a single database. Typically, oral commensal producers were represented by the genera *Capnocytophaga*, *Fusobacterium*, *Haemophilus*, *Neisseria* and *Prevotella*. Strikingly, 94% of hits came from allochthonous microorganisms (*e.g.*, diverse <u>e</u>nteric <u>G</u>ram-<u>n</u>egative rods – EGNRs, and staphylococci) raising the question about their prevalence in human oral cavity and their clinical relevance.

We aggregated clinical findings from 57 studies encompassing more than 27,000 patients from 14 countries and multiple clinical conditions to show that EGNRs are prevalent in human oral cavity [median = 22%, mean = 29% ± 6% (95% CI), **Fig. 3a**]. The prevalence of EGNRs appeared higher in pathologic conditions (ANOVA F = 3.8, *P* = 0.016, **Fig. S4a**), slightly increased in adults (**Fig. S4b**), differed between countries (ANOVA F = 4.6, *P* = 0.0029, Tukey’s Adjusted *P* = 0.019 – 0.064 for 4 comparisons, **Fig. S4c**), and was potentially overestimated in small studies (ANOVA F = 2.2, *P* = 0.12, **Fig. S4d**). Taxonomic data identified *Enterobacter cloacae*, *Escherichia coli*, *Klebsiella pneumoniae*, *Pseudomonas aeruginosa* and *Staphylococcus* species as most prevalent allochthonous species across studies (**Fig. 3b**). These microorganisms were confirmed as significant β-lactamase producers based on genomic information (n = 48,564) from the CARD database^55^. For example, genomes of *Enterobacter cloacae* (n = 244) carried *bla*_CMH-1_ (42%) and *bla*_TEM-1_ (21%), *Escherichia coli* (n = 29,177) carried *bla*_AmpC_ (53 – 66%) and *bla*_TEM-1_ (14%), *Klebsiella pneumoniae* (n = 12,425) carried *bla*_SHV-11_ (48%) and *bla*_TEM-1_ (30%), *Pseudomonas aeruginosa* (n = 4,777) carried *bla*_OXA-50_ (74%) and *Staphylococcus* species (n = 1,941) carried *blaZ* (65 – 85%). It should be noted that these strains represented diverse clinical origins and may not fully reflect the characteristics of oral isolates.

**Fig. 3.**
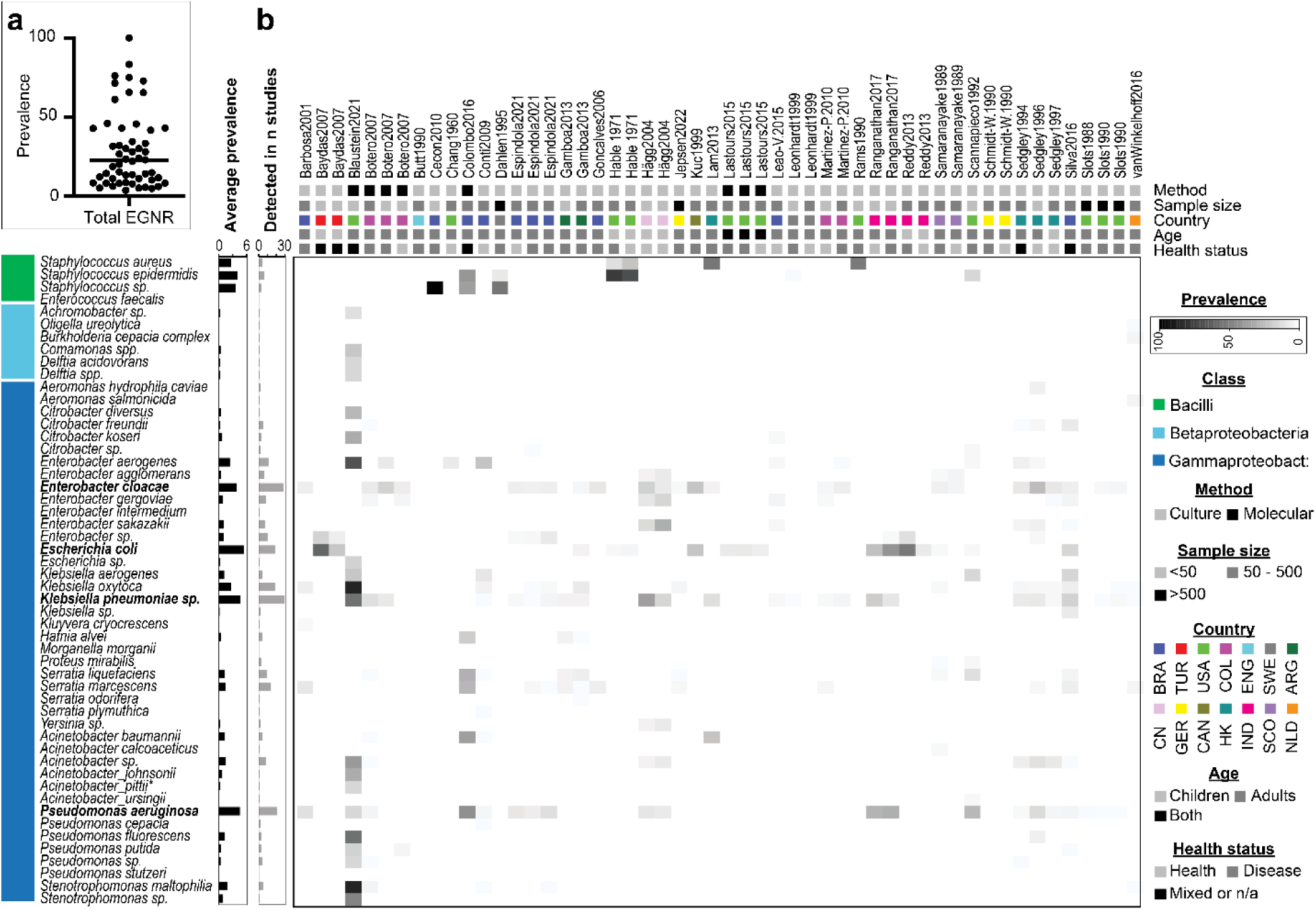
Prevalence of selected oral allochthonous species. **a** Prevalence of oral enteric Gram-negative rods in 27 thousand patients from 57 studies. **b** Taxonomy of selected detected oral allochthonous species across 52 studies.

In summary, multiple lines of evidence confirmed that diverse oral microorganisms (including allochthonous invader species) representing nine classes and more than 80 species have the genetic potential to produce β-lactamases (**Tab. 1**). Four resident (*Porphyromonas*, *Prevotella sensu lato*, *Capnocytophaga* and *Haemophilus*) and five invader genera (*Bacillus*, *Staphylococcus*, *Enterobacter*, *Escherichia*, *Klebsiella* and *Pseudomonas*) were selected for further analyses according to high prevalence in biofilms and common ability of producing β-lactamases. Diverse enzymes representing four major classes were identified. Some of these enzymes can be occasionally produced in large amounts (hyper-produced) if gene expression is high (*e.g.*, due to high gene copy numbers, strong gene promoters, or dysregulated expression control).

**Tab. 1.**
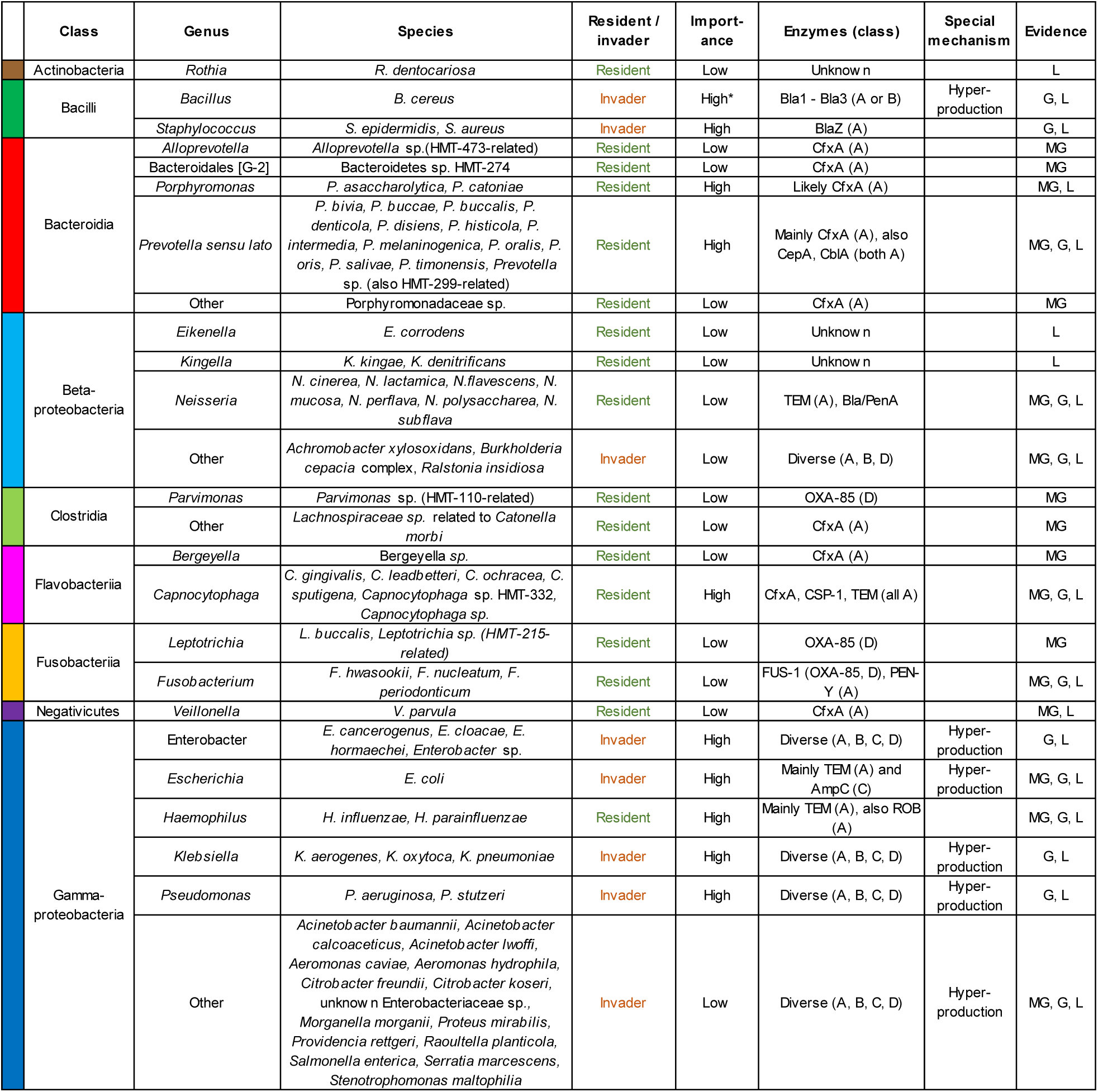
Oral β-lactamase producers. Listed species were sorted by their class and genus taxonomy. Genera were defined as resident or invader with low or high importance. Important genera incorporated species which are both (i) prevalent inhabitants or human oral cavity and (ii) their strains are often able to produce β-lactamases or at least have a genetic potential to do so. Prominent enzymes with their classification and potential for hyper-production as well as lines of evidence were given for each genus. G –<u>g</u>enomes, L – literature, MG – <u>m</u>eta<u>g</u>enomics assemblies (**M&M**). * High importance of *Bacillus* was assigned based on data for specific HIV-positive population and not for general population.

### Multiple allochthonous species and only a single commensal are able to cross-protect

Genetic potential for producing β-lactamases in oral microbes described in a previous section does not necessarily imply that they are able to cross-protect. In order to link the presence of β-lactamase gene with an ecological function, we developed the simple but robust agar diffusion plate assay (**Fig. 4a**). We compared the cross-protection activity of strains (n = 118) representing different species (n = 22), multiple enzymes and several hyper-production mechanisms. 96 cross-protecting strains represented two classes (*i.e.*, Bacilli and γ-Proteobacteria), eight species and mainly four enzymes (BlaZ, TEM, SHV and AmpC) of either A or C class. In these experiments cross-protection was conferred by different invaders (*Staphylococcus*, *Enterobacter*, *Escherichia*, *Klebsiella* and *Pseudomonas* species) and only one commensal genus: *Haemophilus* (**Fig. 4b**, **Fig. S5**). Strikingly, none of the strains representing the class Bacteroidia conferred cross-protection in our test setting.

**Fig. 4.**
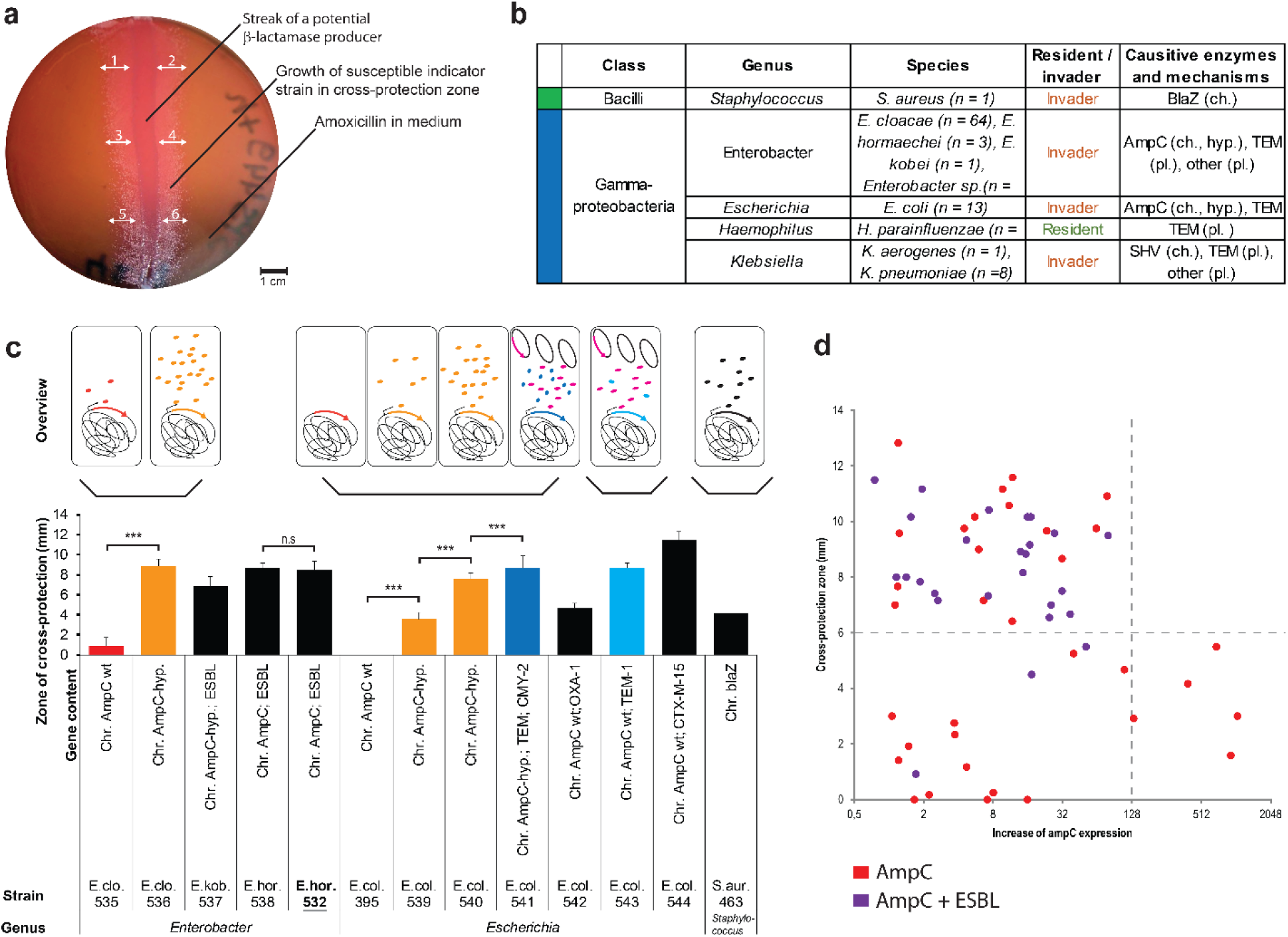
Species conferring cross-protection from Amoxicillin and the major underlying mechanisms. **a** Diffusion agar plate assay for cross-protection between strains. **b** Cross-protecting species. **c** Key cross-protection mechanisms. **d** Cross-protection as an effect of interplay between expression level of chromosomal *bla*_AmpC_ and carriage of ESBL *bla* gene in E. cloaceae strains.

Depending on the Amoxicillin concentration, cross-protection was either non-specific or specific. At a lower concentration of 1.6 µg/mL, virtually all studied susceptible commensal species got cross-protected. At a higher concentration of 8 µg/mL, mostly expansion of *Capnocytophaga* spp. and *Prevotella* spp. was favored.

### Cross-protection activity was often conferred either by a high expression of chromosomal bla_AmpC_ gene or by the presence of plasmidic bla_TEM-1_ gene

Production of TEM by the reference helper strain *Enterobacter* sp. SPS_532 originally led to the detection of polymicrobial resistance (see first section from the **Results**). The discovery that only specific species and strains are able to cross-protect raised the question about further responsible enzymes and specific mechanistic underpinnings. Therefore, we compared the activities across strains producing different enzymes (either chromosomal or plasmidic) at different levels.

Strains carrying chromosomal *bla*_AmpC_^wt^ (*i.e.*, the wild type gene encoding the class C AmpC enzyme) did not cross-protect, irrespective of the studied species and gene inducibility (with the exception of *Pseudomonas aeruginosa*) (**Fig. 4c**, **Fig. S5d**, **Fig. S5e**). In contrast, increased expression, also known as hyperproduction (*i.e.*, presence of *bla*_AmpC_^hyp^) either in clinical isolates (**Fig. 4c**) or mutants (**Fig. S5d**) conferred the activity. A striking example was the gain of activity by *E. cloacae* strain SPS_536 compared to the strain SPS_535 (**Fig. 4c**). Only a single amino acid substitution (Gln86Leu) in *ampD* caused by SNP A256T distinguished the strains SPS_535 and SPS_536 ^56^, but the mutation resulted in a 40-fold increased expression of *bla*_AmpC_ in SPS_536. No differences were found between these isolates in any other gene sequenced. In *E. coli*, cross-protection was induced by the presence of the insertion element ISEc10 (SPS_539) or a point mutation (SPS_540) (**Fig. 4c**). Exposure to Cefpodoxime selected for AmpC hyperproducing *Enterobacter* and *Klebsiella* mutants that exhibited cross-protection activity (**Fig. S5d**). Other chromosomal β-lactamases genes (*bla*_SHV_ and *blaZ*) conferred the activity in *Klebsiella* and *Staphylococcus* strains, respectively (**Fig. 4c**, **Fig. S5f**).

Plasmidic β-lactamases genes coding for TEM contributed to cross-protection activity in multiple strains representing different species (**Fig. 4c**, **Fig. S5**), including the only commensal helper, *Haemophilus parainfuenzae*. Interestingly, for invader species, the presence of multiple *bla*_TEM_ genes (*e.g.*, SPS_648 – 653) and additional hyperproduction of AmpC in TEM producers (*bla*_AmpC_^hyp^, *bla*_TEM_, *e.g.*, SPS_662 – 663) led to increased cross-protection activity in both clinical isolates and mutants (**Fig. 4c**, **Fig. S5d**, **Fig. S5e**). An extreme example was *E. coli* strain SPS_541, in which all the mechanisms acted together. That strain carries critical point mutation inducing AmpC production (like SPS_540) but additionally carries a plasmid with two genes: *bla*_CMY-2_ linked to ISEcp1-like element, as well as, *bla*_TEM_. Other plasmidic β-lactamases (*e.g.*, *bla*_CTX-M-15_, *bla*_OXA_) also conferred or contributed to activity in different species (**Fig. 4c**, **Fig. S5e**, **Fig. S5f**) but are less common^55^.

To evaluate the correlation between expression of *bla*_AmpC_, presence of additional *bla* genes (including those coding for ESBLs), and cross-protection activity we used 65 additional well characterized *E. cloacae* strains^57^ (**Fig. 4d**). We observed three main groups of strains with following characteristics: (i) low expression – no or weak cross-protection, (ii) low to moderate expression – strong cross-protection and (iii) very high expression – moderate cross-protection. Presence of ESBLs was associated with strong cross-protection. Strongly cross-protecting non-ESBL isolates with low to moderate *bla_AmpC_* expression predominantly carried the *bla_TEM_* gene.

In summary, taxonomically diverse strains conferred cross-protection as long as they were potent β-lactamase producers. The common key mechanisms mainly included: (i) hyperproduction of chromosomal AmpC of the C class, and (ii) expression of plasmidic TEM of the A class. The effect of these mechanisms on cross-protection in biofilms was further studied *in vitro* using model *Enterobacter* and *Escherichia* strains.

### Cross-protection in complex biofilms depends on helper population size, which can be controlled with phage treatment

To characterize the impact of cross-protection on multi-species biofilm development in liquid medium, we initially studied a simple approach consisting of exemplary oral biofilm former, *Streptoccocus mutans* SPS_473 and several exemplary *Enterobacter* helper strains [*i.e.*, SPS_532 (*bla*_AmpC_^wt^, *bla*_TEM_), SPS_536 (*bla*_AmpC_^hyp^) with SPS_535 (*bla*_AmpC_^wt^) used as a control] (**Fig. S6**). When *S. mutans* was co-cultured with the *bla*_AmpC_^wt^ and bla_TEM_-expressing strain SPS_532 in the presence of amoxicillin, they formed biofilms as assessed by confocal microscopy. *S. mutans* cells were protected from the effect of amoxicillin, as proved by plating biofilms for colony forming units. The same observation was true for the *bla*_AmpC_-overexpressing SPS_536 but not for the wild type strain SPS_535. Thus, we reproduced cross-protection *via* two main mechanisms and created an experimental basis for development of more complex biofilm models.

To replicate cross-protection in clinically relevant complex microbial communities, we developed a polymicrobial biofilm model (**Fig. 5**, **Fig. S7**). Key components of the model include: (i) a dental plaque or complex defined inoculum ensuring high microbial diversity, (ii) a helper strain producing β-lactamase, (iii) amoxicillin, and (iv) a bacteriophage specifically targeting for the helper strain without affecting other community members (**Fig. 5a**, **Fig. S8**). We monitored planktonic growth, biofilm formation, β-lactamase activity (**Fig. S7**, **Fig. S9**), diffusion agar profiles (Fig. S10), and full 16S rRNA gene amplicon profiles. Controls showed that complex biofilms cannot develop without the helper in the presence of amoxicillin, the helper alone is a poor biofilm former, and purified phage preparations lack β-lactamase activity (**Fig. S7a** – **Fig S7c**).

**Fig. 5.**
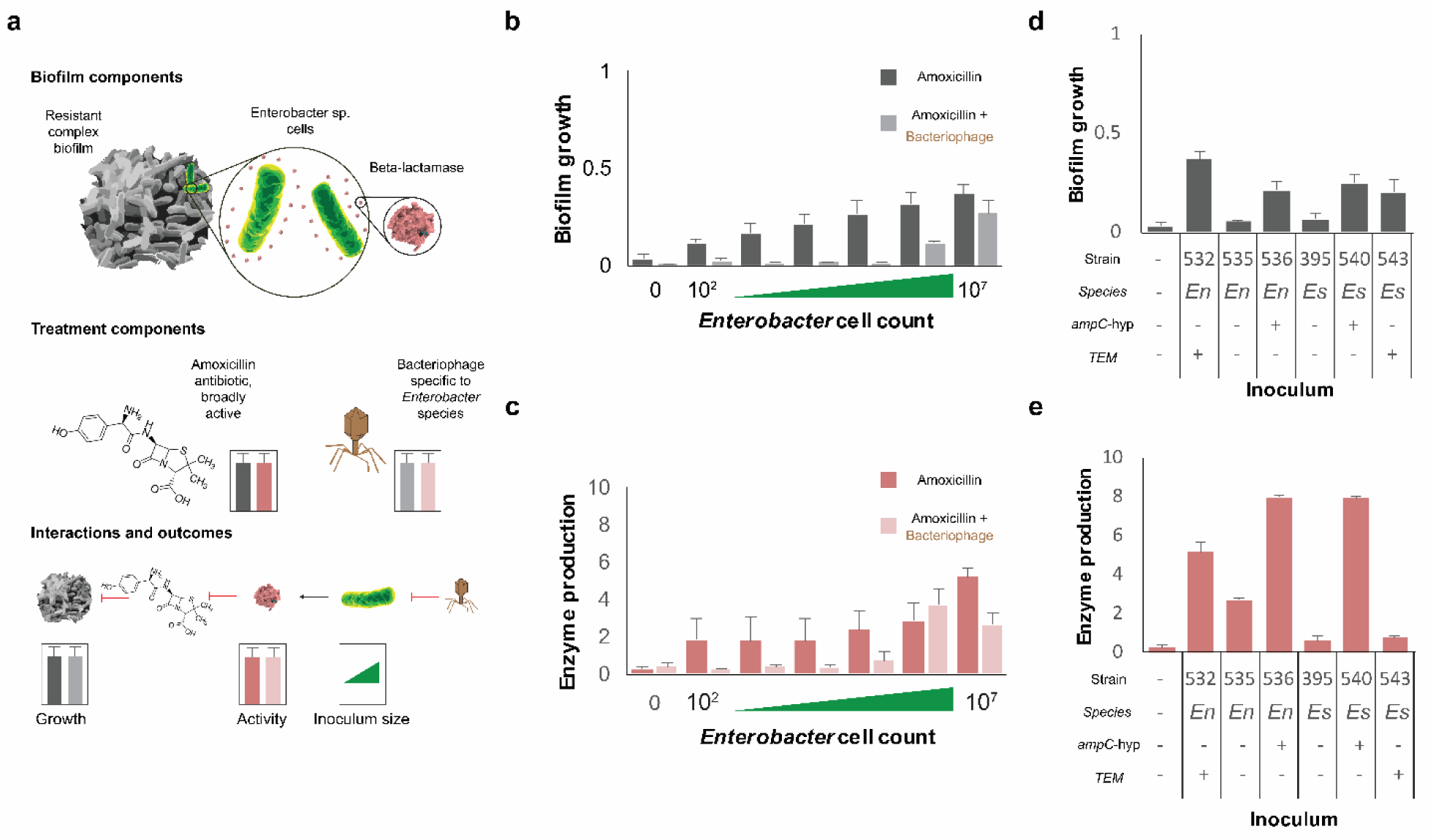
Cross-protection in complex biofilms and its phage-based control. **a** Framework of complex biofilm model. **b** Relationship between complex biofilm growth and helper’s inoculum size in two treatment groups. **c** Relationship between β-lactamase activity and helper’s inoculum size in two treatment groups. **d** Effect of AmpC hyperproduction and TEM production on complex biofilm growth. **e** Effect of AmpC hyperproduction and TEM production on β-lactamase activity. *En* – *<u>En</u>terobacter*, *Es* – *<u>Es</u>cherichia*, hyp – <u>hyp</u>er-production.

The amount of biofilm formation showed a strong positive correlation with the inoculum size of the helper strain (Pearson’s r = 0.65, increasing to r = 1.00 after log_10_ transformation of inoculum size). Even as few as 100 cells of the reference helper strain (*Enterobacter* sp. SPS_532 producing TEM-1) were sufficient to stimulate complex biofilm formation in the presence of amoxicillin (**Fig. 5b**). Phage treatment was effective against inocula containing up to 10^5^ cells, but when phages were omitted, β-lactamase activity remained constant across most inoculum sizes, except for 10^7^ cells where it was higher (**Fig. 5c**). Phage addition reduced β-lactamase activity when the *Enterobacter* inoculum size was below 10^6^ cells.

Interestingly, strong biofilm inoculum-specific effects (*i.e.*, patient-specific effects) were observed for growth, enzyme production and phage susceptibility (**Fig. S7d** – **Fig S7e**). Both hyper-production of AmpC and production of TEM-1 conferred the cross-protection in complex biofilms (**Fig. 5d**). Generally biofilm growth and β-lactamase activity followed the same trend, except for strain SPS_543 (**Fig. 5e**). The variability between inocula was captured (**Fig. 5b** – **5e**, **Fig. S7f**).

### Cross-protection introduced major changes in biofilm composition which were reversed by a phage treatment

We next analyzed the composition of biofilms from the conditions described previously to assess how cross-protection influences the composition of the biofilm. Using an agar diffusion plate assay, we found biofilm tolerance ranged from none to high, with extreme resistance phenotypes shown in (**Fig. 6a**, **Fig S10**). Resistant members, predicted to typically represented by a single species (characterize by a single colony type, **Fig S10**), included isolates such as *Capnocytophaga* and *Prevotella*, along with allochthonous species and fungi, though none displayed cross-protection activity. Cross-streaking with helper strains increased both the number and diversity of colony biofilms, including the emergence of black-pigmented colonies (**Fig. 6a**, **Fig S10**).

**Fig. 6.**
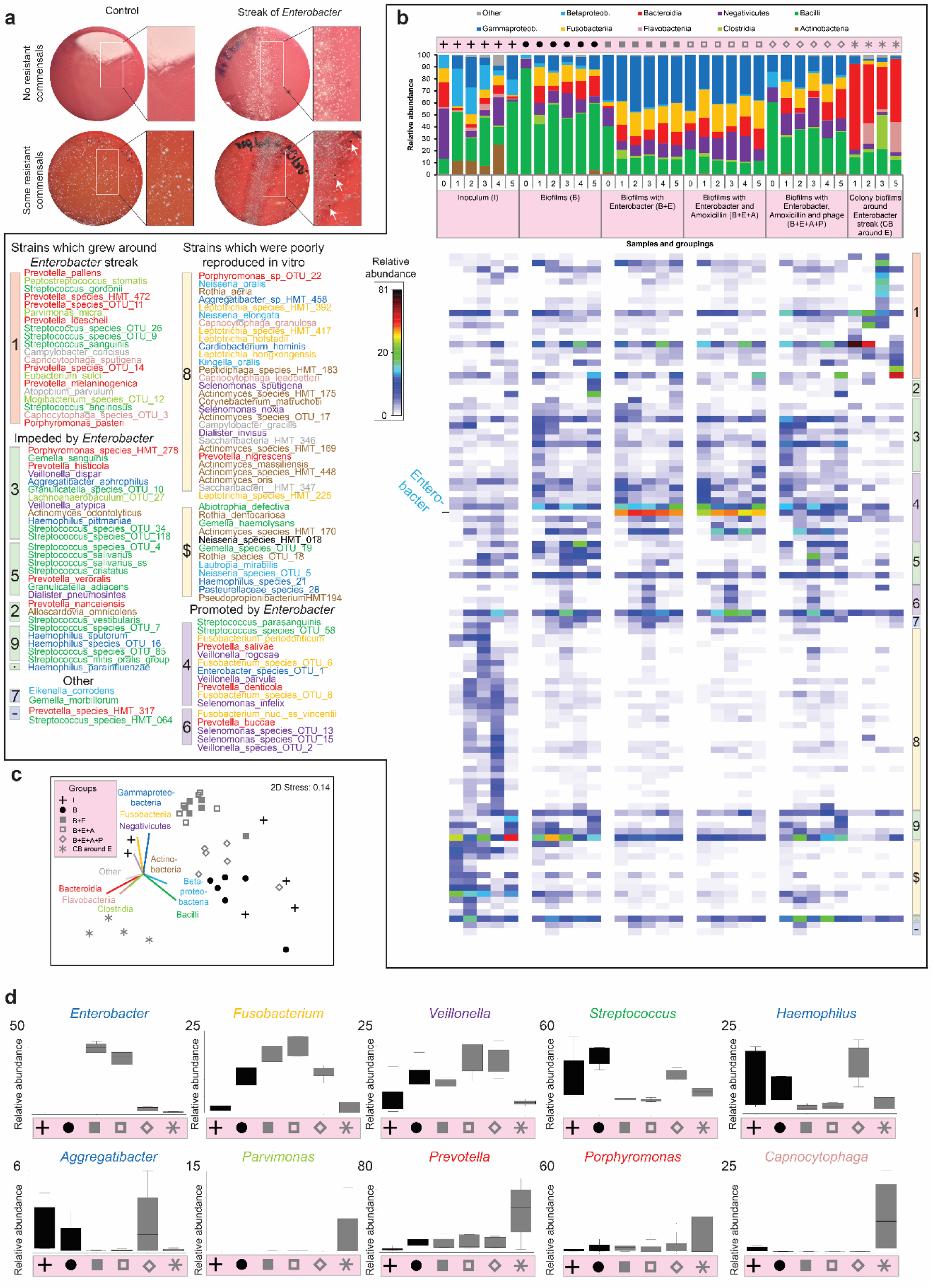
Effect of cross-protection and phage treatment on composition of complex biofilms. **a** Diffusion agar plate assay for cross-protection between a helper strains and complex biofilms. **b** Composition of complex biofilms at class and species level in six experimental groups. The groups consist of clinical inocula (I), *in vitro* biofilms (B), biofilms supplemented with *Enterobacter* helper (B+E), and additionally with Amoxicillin (B+E+A), and additionally with *Enterobacter* phage (B+E+A+P) as well as satellite colony biofilms formed on medium with Amoxicillin around *Enterobacter* streak in diffusion assay (CB). Biofilms were inoculated by a defined and well-characterized collection of 19 strains from a single individual with peri-implantits (0) or supragingival plaque samples from healthy donors (1 – 5). **c** Ordination visualizing the relationships between biofilm compositions at the class level for different experimental groups. Vector overlay indicating the major microbial classes in biofilms. **d** Effect of cross-protection and phage treatment on composition of complex biofilms; a summary. Box plots show the relative abundance for the ten most important genera and different experimental groups.

We profiled the composition of complex biofilms in high resolution and depth using PacBio SMRT sequencing of full 16S rRNA gene amplicons (**Fig. 6b – 6d, Fig. S11**). This yielded a total of 560,000 long reads, averaging 16,500 reads per biofilm (**Fig. S11a**). The experimental groups included inocula (I), *in vitro* biofilms (B), biofilms supplemented with the *Enterobacter* helper strain (B+E), biofilms with both *Enterobacter* and amoxicillin (B+E+A), and biofilms further supplemented with *Enterobacter* phage (B+E+A+P). Additionally, we analyzed satellite colony biofilms formed on amoxicillin-containing media surrounding *Enterobacter* streaks in the diffusion assay (CB) (**Fig. 6b**). All biofilms were inoculated with a defined and well-characterized collection of 19 strains sourced from a single individual with peri-implantitis (0) or supragingival plaque samples from healthy donors (1–5).

Major shifts in biofilm composition were observed between experimental groups, displayed at a broader class (**Fig. 6b**, bar plots) and genus (**Fig. 6d**, box plots) level for clarity and as a vector overlay on the ordination (**Fig. 6c**). Compared to the inoculum, Bacilli were enriched *in vitro* at the expense of Actinobacteria and β-Proteobacteria, while biofilms remained highly diverse, except for biofilms “0”, which were formed by 19 defined strains collected from a single patient (**Fig. S11**). In groups containing *Enterobacter*, γ-Proteobacteria expanded regardless of the presence or absence of amoxicillin. In these groups, Fusobacteria also increased, while Bacilli abundance decreased. The addition of phages (MOI = 1.6 × 10⁴) reversed these trends. In contrast, entirely different microbial communities were observed in the diffusion assay, which did not resemble any other experimental group and were dominated by Bacteroidia, Flavobacteriia, and Clostridia.

At the species level (**Fig. 6b**, shaded plot), we observed distinct microbial patterns specific to both the inocula (PERMANOVA Pseud-F = 7.6, *P* < 0.0001; pairwise t > 1.4, *P* < 0.05 in 8 out of 10 comparisons) and the experimental groups (PERMANOVA Pseud-F = 6.0, *P* < 0.0001; pairwise t > 1.77, P < 0.01 in 13 out of 15 comparisons) (**Fig. S11e**). The inocula were characterized by a dominance of commensals with low pathogenic potential, representing what is considered healthy human oral flora (**Fig. S11d**). We successfully reproduced 80% of the species richness *in vitro*, with an average richness of 133 compared to 164 in the inoculum. Biofilms inoculated with a defined strain collection (across all experimental groups), as well as colony biofilms from the diffusion assays, exhibited significantly lower diversity (compared to biofilms inoculated by complex dental plaques and cultured in liquid), as anticipated (**Fig. S11**). Biofilms containing *Enterobacter*, regardless of amoxicillin presence, were similar in composition. Interestingly, phage-treated biofilms resembled those without *Enterobacter*, effectively reversing the influence of *Enterobacter* on biofilm composition.

## Discussion

The effective management of polymicrobial biofilm infections may be improved by targeted antimicrobial therapies to eradicate species that are central to the fitness and survival of the entire biofilm as demonstrated in multiple studies on the gut microbiome^58, 59^. However, in oral microbiology, such approaches have historically been underappreciated, likely due to methodological challenges in their development for complex biofilms and a lack of convincing evidence demonstrating their significance.

In this study, we adapted laser-based bioprinting for antimicrobial susceptibility testing of complex biofilms and observed polymicrobial β-lactamase-driven amoxicillin resistance in peri-implantitis. By leveraging genomic (n = 200,000) and microbiological data (n = 27,000), a representative strain collection (n = 118), cross-protection assays and a complex biofilm model (average 133 species), we showed that this phenomenon is primarily caused by the few allochthonous species mostly releasing either AmpC or TEM-1 β-lactamases. Using the dental plaque as a showcase example, we demonstrated that even a single point mutation in the genome of a β-lactamase producer could initiate the development of cross-protection which not only shielded the biofilm as a whole but also promoted the expansion of opportunistic pathogens. By using a phage isolate specifically targeting the β-lactamase producer for elimination, we confirmed that cross-protection could be effectively bio-controlled. However, the efficacy of this intervention depended on the population size of the producer, underscoring the importance of timely application.

Recent studies have shown that antibiotic resistance can arise as a result of combined effects of antibiotic exposure and microbial interactions within communities^13^. However, these interspecies interactions are overlooked when antibiotic resistance is investigated in monocultures. Microbial arrays created with laser-assisted bioprinting allow diffusion of metabolites and enzymes between microbial colonies and in this way can preserve some of the *in vivo* interactions^18^ making it a valuable method to detect polymicrobial resistance. The phenomenon of protective clearance of antibiotics by resistant cells is commonly seen by microbiologists in the presence of ‘satellite’ colonies on plates with clinical materials or transformation plates. But in such case, in contrast to bioprinted microbial arrays, the position of colonies is random and unordered^60^. Moreover, due to its technical parameters and modular structure, bioprint technology serves as a strong foundation for further development, such as integration with high-throughput sequencing or hybridization techniques. Our data from bacterial mutants indicate that strong cross-protection can be readily induced in diverse members of the Enterobacteriales order by expanded-spectrum cephalosporins. This finding aligns with existing literature^61, 62^ and highlights the need for easy-to-perform and accurate diagnostics which support the clinician’s management decision for patients carrying such strains. Furthermore, distinguishing between TEM-1 and AmpC-based cross-protection is crucial, as these enzymes exhibit different susceptibility profiles to inhibitors^63^.

Historically, clinical data, animal experiments, and mathematical models have suggested that β-lactamase-driven cross-protection is a relevant resistance mechanism in polymicrobial infections across various body sites^22, 43, 64–68^. However, this phenomenon has not been systematically studied in context of the oral microbiome so far. Our extensive study, incorporating data from a large number of patients and strains, identified allochthonous Gram-negative *Enterobacter*, *Escherichia* and *Klebsiella* species as well as Gram-positive staphylococci as key drivers of cross-protection. *Enterobacter* and *Klebsiella* species are metabolically adapted to thriving in an inflammatory milieu but can also survive well in home environments, *e.g.*, on surfaces or toothbrushes^25, 49, 50^. They show a great ability to be transmitted, which weakens the selection for mutualistic relationship with the host^69^. Poor oral hygiene, hospitalization, inadequate household hygiene, and systemic antibiotic treatment promote oral colonization^70^, with the latter disrupting colonization resistance conferred by activities of indigenous oral flora^71, 72^.

*Enterobacter* and *Klebsiella* species harbor β-lactamase genes chromosomally or additionally on plasmids and are intrinsically resistant to amoxicillin. Exposure to β-lactam antibiotics like cephalosporins and carbapenems, used to treat severe infections, can drive resistance further by upregulated expression of resistance traits or acquired resistance genes and their accumulation^73^. We found that both mechanisms can contribute to cross-protection activity and act synergistically, *e.g.*, in strains harboring derepressed *bla_AmpC_* and *bla_TEM_* or expended spectrum β-lactamase bla_CTX-M-15_ genes on plasmids.

Our findings are of broad significance beyond the dental field. The oral reservoir of allochthonous β-lactamase producers in susceptible patients, including the elderly, disabled, immune compromised and hospitalized individuals^74–77^, may significantly impact overall health by enabling the systemic dissemination of these organisms. This can lead to their migration to the gut, potentially exacerbating inflammation^78^, or to the airways, where they may cause pneumonia^76, 79^. Therefore, tailored hygiene plans should be implemented for these patients to mitigate these risks.

*Enterobacter* and *Klebsiella* species exhibit remarkable resilience to starvation, mechanical therapy and treatments with silver and chlorhexidine, making their eradication challenging^46, 47, 49, 50^. They possess intrinsic resistance to aminopenicillins like amoxicillin and frequently (79% of *Enterobacter* sp. strains) or occasionally (5 – 6%, *Klebsiella*) resist combination of amoxicillin and clavulanic acid and doxycycline (20% and 6 – 9%, respectively)^80^. Additionally, they exhibit occasional resistance to ciprofloxacin (1.5 and 2%)^80^ although these numbers vary across different populations^45^. These species are often key contributors to severe, treatment-resistant infections, regardless of the drug combinations used^81^. *In vitro* sensitivity data suggest that therapy combining ciprofloxacin and metronidazole could promote a subgingival microbiota predominantly composed of beneficial streptococci^45^. Similarly, we achieved a comparable and promising therapeutic outcome in our complex biofilm model by combining a lowered concentration of amoxicillin with phage treatment targeting cross-protecting strains. This supports the potential of phage-based anti-biofilm strategies for lowering the risk of post-treatment emergence of superinfection and clinical treatment failure^82, 83^. Recent proof-of-concept studies on *Klebsiella*-targeting phages in the context of IBD^84^ further highlight their potential, suggesting that adjunctive oral phage therapy could be a viable option. A deeper understanding of colonization resistance is crucial for developing pre- and probiotic-based strategies to prevent the re-colonization of *Enterobacter* and *Klebsiella* species^53, 71, 85, 86^.

*Haemophilus* species were the only indigenous oral bacteria that provided cross-protection in our experimental setting, consistent with previous reports identifying *Haemophilus* producer strains in clinical samples^87^. In contrast to a previous animal study^43^, we were not able to identify indigenous *Prevotella* helper strains. This discrepancy may be due to differences in antibiotic concentrations used or the absence of relevant *Prevotella* strains in our collection or studied clinical samples.

Our complex *in vitro* biofilm experiments, along with the plate diffusion assay, revealed that β-lactamase producers facilitate the growth of certain opportunistic pathogens, such as Fusobacteria and Bacteroidia. Consequently, cross-protection not only shields polymicrobial biofilms but also enhances their diversity and virulence. This aligns with findings from *in vitro* experiments using specific genetic constructs, which demonstrated that traits benefiting other group members, specifically sharing β-lactamases, can enable the survival of community members that would otherwise be eliminated by natural selection, thereby maintaining greater genetic variation^88^ and virulence^89^. However, the beneficial effects of β-lactamase producers have also been observed, particularly in supporting the recovery of natural gut flora after antibiotic treatment^90^.

It has been hypothesized that phenotypically susceptible bacteria might rarely benefit from detoxification by others, whereas persisters are more likely to exploit the β-lactamases of their neighbors^60^. We found that this effect is diffusion-dependent, as expansion of more tolerant species was observed on diffusion assay plates but not in the complex biofilm cultured in liquid medium. In this case, *Capnocytophaga* and *Prevotella* species survive the initial high concentration of amoxicillin by their own production of β-lactamases, but crucially their ability to grow in our experimental setting is a social trait as it is dependent on the presence of neighboring β-lactamase producers. Basal β-lactamase production, therefore, facilitates the exploitation of antibiotic-reduced space provided by the helper cells. This specific effect was not observed in the complex biofilm system, which included liquid medium overlaying the biofilm structures, and which was characterized by much greater exchange of metabolites and universal cross-protection, in agreement with data from similar previous studies^91^.

Our experimental settings are not without limitations, highlighting the need for further research. Implementing medium exchange or using a flow chamber model could better replicate the conditions observed during clinical treatment of bacterial infections^91^. Additionally, metatranscriptomic analysis could reveal microbial adaptation mechanisms and clarify how *Enterobacter* sp. facilitates the expansion of Fusobacteriia. Expanding the range of experimental parameters, including strain diversity, concentrations, duration, and mode of intervention, through both *in silico* and *in vitro* studies would provide a more comprehensive understanding of the phenomenon. From a clinical perspective, studying the impact of various treatments on the emergence of cross-protection would be valuable^81^. Additionally, assessing the relevance of these findings across diverse populations could be achieved through a multinational study on the prevalence of cross-protection, its contribution to disease and correlation with treatment failures. The potentially relevant patient demographic likely includes individuals with periodontal or peri-implant disease who are non-responsive to β-lactam antibiotics and have a history of second-generation cephalosporin treatment. In our study we have not clarified how the β-lactamases are released but the broad taxonomic diversity of cross-protecting strains suggest multiple potential mechanisms, including leakage from viable cells of Gram-positive bacteria, release from lysed or broken cells of Gram-negative cells or exported as cargo of membrane vesicles^92^. Factors influencing release mechanisms and consequently cross-protection activity require further investigations.

As a step toward further personalized medicine and to meet the demand for alternative strategies against biofilm-infections, we highlighted the crucial role of bacteriophages, which are highly selective and biologically accepted but have not been sufficiently studied in dental medicine. We unveiled their potential in regulating bacterial populations in various ecosystems. Making their comeback into western medicine, phages are fascinating entities that hold immense potential for therapeutic applications. We demonstrated that particularly in treating antibiotic-resistant infections they combat specific strains while minimizing harm to beneficial bacteria.

In summary, our data suggest that cross-protection involving TEM and AmpC β-lactamases may play a significant role in the polymicrobial resistance of oral biofilms to amoxicillin. Allochthonous *Enterobacter* and *Klebsiella* species may have a more important ecological role in oral biofilms than previously thought. Diagnostics, potentially involving laser-based methods, and therapeutics tailored to addressing cross-protection and allochthonous species could improve patient management, particularly in vulnerable populations.

## Materials and methods

### Collection of clinical samples

This study analyzed samples from the SIIRI Peri-Implant Biofilm Cohort, a long-term initiative aimed at monitoring the peri-implant microbiome over a period of up to 12 years to investigate disease development and prevention. The study was conducted at the Department of Prosthetic Dentistry and Biomedical Materials Science, Hannover Medical School, Germany, and was approved by the institutional ethics committee (approval numbers 5544 and 9477). Written informed consent was obtained from all participants. Peri-implantitis was diagnosed according to the case definitions given by the World Workshop on the Classification of Periodontal and Peri-implant Diseases and Conditions, 2017^93^. Clinical examination was performed as previously described^94^. For microbiological analysis, biofilms associated with submucosal implants were collected using a curette, transported in reduced transport fluid (RTF)^95^ and further processed as previously described^18^.

### Antimicrobial susceptibility testing of peri-implantitis biofilms using laser-assisted bioprinting

Three separate oral biofilm samples from peri-implantitis sites were vortexed in 500 µl of RTF and aliquoted: up to 40 µl of fresh material for laser-assisted bioprinting, 50–100 µl with BHI-glycerol for storage and future printing or plating, 50–100 µl for DNA isolation, 70–200 µl for microscopy, 150 µl for classical plating, and any remaining volume, if applicable, for additional glycerol stocks. Bioprinting was performed like previously described^18^. Briefly, fragmented biofilms were suspended in a bioink consisting of 4 parts human fibrinogen (40mg/ml), two parts glycerol and one part hyaluronic acid (1% in TBS, all from Sigma Aldrich) and placed on a donor glass slide as a bioink layer. This bioink was printed as droplets, each of which was propelled from this layer by one laser pulse, on solid media with antibiotics in form of classical petri plates, or on square agar slices with a length of 23 mm, height of 5 mm, and an antibiotic disc placed in the center. Up to 440 colony-like biofilms we printed in 1 mm distances, forming a growth pattern of dots in a square grid on the cultivation cubes. Antimicrobial susceptibility testing of biofilms was performed anaerobically (80% N_2_, 10% H_2_, and 10% CO_2_) using enriched Brucella blood agar, or aerobically (5% CO_2_) using enriched Mueller Hinton Agar. In both cases, antibiotics were either incorporated directly into the medium or applied via antibiotic-impregnated disks. The first basis medium (MSPS_042) contained 43.1 g/L Brucella Agar (5752.1, Carl Roth), 5% defibrinated sheep blood, 5 mg/L haemin, 5 mg/L L-histidine, and 10 mg/L vitamin K_1_. The second basis medium (MSPS_048) contained Mueller-Hinton Agar (CM0405, Oxoid), supplemented with 17 g/L agar, 20 mg/L NAD and 5% defibrinated sheep blood. Microorganisms resistant or tolerant to specific antibiotics were isolated on those basis media supplemented with antibiotics: 8 mg/L amoxicillin (A8523-1G, Sigma-Aldrich, MSPS_042A and MSPS_048A), 16 mg/L metronidazole (M1547, Sigma, MSPS_042B and MSPS_048B), 8 mg/L tetracycline (HP63.1 Roth, MSPS_042C and MSPS_048C) and 0.5 mg/L ciprofloxacin (17850-5G-F, Sigma-Aldrich, MSPS_042F and MSPS_048F). The breakpoint concentration values for amoxicillin, metronidazole, tetracycline and ciprofloxacin are based on previous studies of periodontal and dental implant microbiology^34, 96, 97^ and reflect drug levels at or above the minimal inhibitory concentrations associated with the “susceptible” interpretative category for anaerobic bacteria as defined by the Clinical and Laboratory Standards Institute (CLSI M11 9^th^ ed.). Plates were incubated aerobically and anaerobically for up to 3 days and evaluated after up to 7 days. A lower concentration of Amoxicillin was also tested (1.6 mg/L, medium designated MSPS_042A2).

Further antimicrobial susceptibility testing was performed either as described above but by using a range of different antibiotic concentrations diluted in the agar, or by using antibiotic disks placed on both basis media. Briefly, we applied the colony suspension method (CLSI M11 9^th^ ed.) to inoculate plates. Bacterial biomass of isolates was resuspended in 0.85% NaCl, with the optical density adjusted to 0.169, corresponding to a McFarland Standard of 0.5. A 50 µL aliquot of each inoculum was evenly spread on either MSPS_042 or MSPS_048 plates. Up to four antibiotic disks were placed on the plates using sterile forceps, and the plates were incubated at 37°C in 5% CO₂ for 24 hours.

### Characterization of β-lactamase producers using (meta)genomics

To describe the spectrum of producers and enzymes relevant for the human oral cavity we combined the following resources: publically available metagenomic^54^ and genomic information^98^. Briefly, we analyzed more than 150 thousand microbial genomes constructed via single-sample assembly from almost 10,000 thousand metagenomes representing diverse human populations and subsequently focused on subsets representing oral representatives^54^. We supplemented this data with more than 2,000 thousand genome sequences from the curated eHOMD database for oral microorganisms^98^. Finally, we characterized the prevalences of β-lactamase genes for oral allochthonous species using information stored in the CARD database of antibiotic resistance determinants^55^. A multi-database search for β-lactamase genes was performed with ABRicate employing NCBI, CARD, ARG-ANNOT, Resfinder, MEGARES, EcOH, PlasmidFinder, Ecoli_VF and VFDB databases (https://github.com/tseemann/abricate).

### Aggregation of literature information on prevalence of selected oral allochthonous species

A comprehensive search was conducted in PubMed using the terms “Enterobacter”, “Escherichia”, “Klebsiella”, “Pseudomonas”, “Serratia”, “Staphylococcus”, “Bacillus”, “enteric rods”, “pseudomonads”, “gram-negative rods”, “Enterobacteriaceae”, “Pseudomonadaceae”, “staphylococci”, or “coliform” in combination with “dental”, “periodontal”, “tongue” or “mouth”. The articles with full-text availability, published in 1975 or later, were retrieved. Titles and abstracts were manually screened for eligibility. The inclusion criterion was clinical studies or review articles addressing the prevalence of oral invader species. References cited in review articles were used to cross-validate the primary search results. Additionally, a secondary validation was performed by manually searching for articles that cited the most highly referenced articles identified in the primary search.

### Validation of genomic predictions with literature information on typical oral species β-lactamase producers

A comprehensive search was conducted in PubMed using a strategy similar to that used for identifying studies on the prevalence of oral invader species. We used terms including genus name, *e.g.*, “Capnocytophaga”, in combination of “lactamase”, *e.g.*, (Capnocytophaga[Title/Abstract]) AND (lactamase[Title/Abstract]). The inclusion criterion was research studies or review articles addressing the detection of lactamases (genes, proteins or activity) in typical oral species.

### Isolation, storage and characterization of clinical strains

To study the cross-protection *in vitro*, we analyzed reference strain collection^18^ expended by additional published and own isolates^57, 99^. Isolation and characterization of strains was performed like described before^18^. Briefly, interesting colonies identified during antibiotic susceptibility testing or colony-like biofilms from biofilm printing were transferred on fresh media. Clones showing unique colony and/or cell morphology were purified through at least three passages and stored in Brain Heart Infusion broth supplemented with 50% glycerol at -80°C. PCR targeting the 16S rRNA genes and Sanger sequencing were used to assign the taxonomy.

### Detection of β-lactamase activity in bacterial spent medium

A colony of SPS_532 was inoculated into 40 mL of THBY medium for liquid culture. After 24 hours of growth, the culture was centrifuged at 4500 × g for 15 minutes at 4°C, and the supernatant was subsequently filtered through a 0.22 µm filter (Carl Roth GmbH+Co KG, Karlsruhe, Germany). The filtered supernatant was divided into four aliquots, each containing 1000 µL. The first aliquot was heated to 95°C for 30 minutes. The second aliquot was adjusted from originally pH 6.5 to pH 7.7, matching the pH of the original THBY medium. The third aliquot, to which Proteinase K (#19134, Qiagen, Hilden, Germany) was added to a final concentration of 200 µg/mL, and the mixture was incubated at 50°C for 45 minutes. After cooling, 10 µL of Protease Inhibitor Cocktail (Halt Protease Inhibitor Cocktail, Thermo Fisher Scientific, Waltham, MA, USA) was added and the sample was vortexed. An additional aliquot of untreated supernatant was kept as control. All aliquots were then stored at 4°C for 24 hours.

Eight plates of MSPS_42A and one plate of MSPS_42 (as a control) were prepared and stored at 4°C for 24 hours. Following this, 50 µL of indicator strain inoculum (*Lactobacillus* sp. SPS_449 in 0.85% NaCl, OD 0.169 McFarland standard) were evenly distributed across each plate. One plate of MSPS_42A was retained as a control, while the remaining plates had holes punched into the center of the agar. Each hole was filled with approximately 500 µL of one of the supernatant aliquots, and the plates were incubated for 48 hours under anaerobic conditions (80% N_2_, 10% H_2_, and 10% CO_2_). The plates were examined visually for the presence or absence of satellite growth around the holes and photo-documented.

### Disc diffusion tests for β-lactamase activity

Disc diffusion assays were performed using combination disc sets D63C, D68C, and D69C (Mast Group Ltd., Merseyside, United Kingdom), following the manufacturer’s instructions. Sets D69C and D68C were employed to detect AmpC-producing Enterobacteriaceae. These tests enable selective detection by combining an AmpC inducer with both ESBL and AmpC inhibitors. Additionally, set D63C was used to confirm ESBL production in Enterobacteriaceae with chromosomal AmpC.

Fresh 24-hour-old bacterial cultures were suspended in Dulbecco’s Phosphate Buffered Saline (Sigma Aldrich, St. Louis, Missouri, USA) to a concentration equivalent to a 0.5 McFarland standard. A 50 µL aliquot of each suspension was evenly spread across a blood agar plate (Mueller-Hinton Agar supplemented with defibrinated blood). Using sterile forceps, the antibiotic discs were placed onto the inoculated medium, ensuring adequate spacing between the discs. Following incubation at 37°C for 18-24 hours, the plates were examined visually to measure the zone diameters around the discs, with results compared for differences.

### Genotypical and functional characterization of β-lactamase producers

The presence of β-lactamase genes (coding for CfxA, CepA/CblA, BlaZ, TEM, SHV, OXA, CTX-M, ACC, FOX, MOX, CMY, DHA, LAT, BIL, CMY, ACT, MIR, GES, PER, VEB, OXA, IMP, VIM, KPC) in the strains was investigated by classical or multiplex PCR^38, 100, 101^. β-lactamase activity was assessed using a colorimetric assay with nitrocefin (#484400, Sigma-Aldrich). The hydrolysis of nitrocefin by β-lactamase, resulting in a color change from yellow to red, was monitored using a microplate reader to measure the absorbance at 490 nm. Additionally, nitrocefin-impregnated disk (Thermo Scientific™ Remel™) were used for rapid detection of β-lactamase production by *Haemophilus*, *Neisseria*, *Staphylococcus* and diverse anaerobic species.

### Cross-protection streak assay

The cross-protection assay was conducted by streaking strains with potential protective properties onto Supplemented Brucella Agar with Amoxicillin, designated as MSPS_042A2 medium. This medium was prepared by dissolving 28 g/L of Brucella broth powder (Becton Dickinson, MD, USA, BBL, REF 211088), which includes 10 g of pancreatic digest of casein, 10 g of peptic digest of animal tissue, 1 g of dextrose, 2 g of yeast extract, 5 g of sodium chloride, and 0.1 g of sodium bisulfite. The medium was further supplemented with 5 mg/L of haemin (Sigma Aldrich, St. Louis, MO, USA, H5533), 5 mg/L of L-histidine (Roth, 3852.1), 10 mg/L of vitamin K1, 5% defibrinated sheep blood, 1.6 mg/L of amoxicillin (Sigma Aldrich, St. Louis, MO, USA), and 15 g/L of agar. Haemin-histidine in water^102^ and vitamin K1 in ethanol solutions, defibrinated blood, and freshly prepared amoxicillin solution (Sigma Aldrich, St. Louis, MO, USA) in 0.1 M phosphate buffer (pH 6, following CLSI M11 guidelines) were added to the cooled, autoclaved Brucella broth with agar. Plates were prepared at least 72 hours prior to experimentation and stored at 4°C. Before use, the plates were briefly left open to dry and allowed to equilibrate to room temperature. The colony suspension method (0.5 McFarland, following CLSI M11 guidelines) was employed to prepare inocula for both indicator strains (referred to as ‘indicators’) and strains with potential protective properties (referred to as ‘helpers’). Well-isolated 24 to 74 hours old colonies (exhibiting similar morphologies) of indicators and helpers were harvested from growth on solid media, typically Fastidious Anaerobe Agar supplemented with 5% defibrinated sheep blood (MSPS_029 medium), and were. Well-isolated colonies and suspended in 0.85% NaCl solution to achieve a turbidity corresponding to a 0.5 McFarland Standard (Carl Roth GmbH + Co. KG, Karlsruhe, Germany, 1307.1), which equates to an optical density of 0.169 at 600 nm, as measured using a Biophotometer D30 (Eppendorf AG, Hamburg, Germany). Fifty microliters of the indicator strain suspension were evenly spread across the surface of each test plate containing MSPS_042A2 medium. Subsequently, 10 µL of the helper strain suspension was streaked in a single line across the entire diameter of the plate using an inoculation loop. The plates were then incubated under anaerobic conditions at 37°C for 48 hours, unless otherwise specified. The reference indicator and helper strains used were *Lactobacillus* sp. SPS_449 and *Enterobacter* sp. strain SPS_532. Indicators and helpers were always plated on medium without antibiotic as a growth control and their cell counts were controlled by plating for colony forming units. *Haemophilus* sp. helper strains were also studied in aerobic atmosphere with enriched CO_2_ to 5%. Plates were examined visually to measure the protection zone diameters next to streak of cross-protecting strain at six fixed positions (**Fig. 4a**).

### Generation of mutants with derepressed *bla_ampC_* gene

In certain members of the Enterobacterales order, the *bla_ampC_* gene, which encodes β-lactamases of class C, is located on the chromosome. In wild-type strains, AmpC expression is typically low but can be induced^61^. Certain β-lactams, such as cefotaxime, can select for variants exhibiting high-level AmpC expression and increased clinical resistance. Mutants with derepressed ampC expression were generated from strains representing *Enterobacter, Citrobacter, Hafnia alvei, Providencia rettgeri, Providencia stuartii, Serratia marcescens*, and *Morganella morganii*. To create these mutants, parallel cultures were prepared for each strain, with 100 µL of approximately 10^5^ CFU/mL bacterial suspension in Mueller– Hinton Broth (Oxoid, Thermo Fisher Scientific, Waltham, MA, USA), and incubated overnight at 37°C in an atmosphere enriched with 5% CO_2_. The cultures were then spread onto selective Mueller–Hinton Agar (Thermo Fisher Scientific, Waltham, MA, USA) containing cefotaxime (8 mg/L; Oxoid) and incubated overnight at 37°C under the same conditions. Following incubation, colonies were passaged three times to obtain pure strains. Derepression of *bla_ampC_* was confirmed using a disk-based assay and its impact on cross-protection was evaluated. Additional mutants were identified as resistant colonies as they were found within the inhibition zones surrounding antibiotic discs that stimulate high-level AmpC expression.

### Phage isolation and cultivation

Bacteriophages targeting the reference helper strain *Enterobacter* sp. SPS_532 were isolated from sewage samples collected at a municipal sewage treatment plant in Hannover, Germany. Phage enrichment and screening were performed using a plate overlay assay. The culture conditions included a one-day incubation in semi-solid LB medium with helper cells (1:30 dilution of overnight culture) under aerobic conditions, with or without 5% carbon dioxide enrichment, at 37°C. Individual plaques were purified through at least three rounds of passaging to ensure strain specificity, resulting in the isolation of four distinct phage strains (SPS_556 – SPS_559). Among these, phage strain SPS_556 displayed clear plaque morphology, the highest titer (8.0 × 10^10 PFU/mL, compared to 4.3 × 10^8 – 1.5 × 10^10 PFU/mL for the other three strains), and the most potent antimicrobial activity against the host strain, making it the preferred candidate for further experimentation. Filtered (0.45 µm) phage samples were purified, concentrated via ultrafiltration (Amicon Ultra-15 centrifugal filter), and stored in SM phage buffer (100 mM NaCl, 8 mM MgSO_4_ · 7 H_2_O, 50 mM Tris-Cl pH 7.5, 0.01% gelatin) at 4°C. Stock viability was monitored using drop spot and overlay assays. Working solutions were prepared using SM buffer without gelatin.

### Cross-protection in dual-species biofilm model

Three-day-old colonies of *Enterobacter sp.* strains (SPS_532, SPS_535 or SPS_536) and *Streptococcus mutans* SPS_473 were grown on THBY agar plates with blood at 37°C in a 5% CO₂ atmosphere, then used to inoculate liquid cultures overnight. Biomass was concentrated by centrifugation, resuspended in Dulbecco’s Phosphate Buffered Saline (Sigma Aldrich, St. Louis, MO, USA) to an OD₆₀₀ of 2.0, and used to inoculate THBY medium with 1% sucrose to an initial OD₆₀₀ of 0.01. Cultures were incubated for 1, 2, or 3 days in 24-well plates with daily medium replacement. Amoxicillin (8 – 512 mg/L) was added at the start or to 16-hour-old biofilms. After incubation, supernatants were removed, biofilms washed, vortexed, and plated to determine colony-forming units.

### Complex model biofilm, cross-protection assay and phage treatment

In the complex biofilm model, cultures were inoculated with fresh dental plaque from healthy individuals. Supragingival biofilms were collected using cotton swabs, transferred into RTF buffer, and disrupted by vortexing. The inoculum was quantified by colony-forming units (CFUs) on FAA agar (anaerobic, 37°C, up to 7 days), colony morphologies were recorded and cross-protection streak assays using *Enterobacter* sp. strain SPS_532 as helper and clinical/culturome samples as indicator were performed. Cultures that grew around the streak were referred to as colonies supported by β-lactamase diffused from the central helper streak, abbreviated as ‘CB’ (for ‘colony biofilms’).

Seven experimental groups were established for each inoculum (I) in liquid cultures: biofilms with inoculum only (B), without or with a helper *Enterobacter/Escherichia* strain (B+E), without or with amoxicillin (B+A and B+E+A) and without or with phage (B+H+P and B+H+A+P). Biofilms were cultured in 96-well plates with 200 µL BHI medium. The final mixture contained 1.2 · 10^6^ ± 8.0 · 10^5^ CFUs of inoculum, 9 · 10^6^ CFUs of helper strain, and 2 · 10^10^ PFUs of phage, with amoxicillin at 8 µg/mL. Plates were incubated at 37°C for 24 hours under anaerobic conditions. Growth was measured for OD_620 nm_ (Tecan Infinite M200 Pro, Tecan Trading AG, Switzerland), and supernatants were collected for a Nitrocefin (NCF (Sigma Aldrich, St. Louis, MO, USA)) assay. Biofilm growth was also assessed for OD_620 nm_ after removing the planktonic phase, while biomass was quantified with crystal violet staining. Briefly, after washing with saline, the biofilms were dried under sterile conditions. Next, 0.1% Crystal Violet was added, followed by incubation at room temperature with gentle shaking at 300 rpm for 15 minutes. The wells were then thoroughly washed with saline three times with and dried under sterile conditions. Absolute ethanol was subsequently added, and the plates were incubated at 300 rpm for four hours at room temperature. The resulting liquid was mixed, transferred to a new 96-well plate, and the optical density was measured at 620 nm.

To investigate the effect of phage on biofilms inoculated with different numbers of helper strain cells (ranging from 10^2^ to 10^7^), and to assess the impact of different helper strains representing various species (either *Escherichia* or *Enterobacter* strains SPS_ 535, 536, 395, 540, 543) and enzymes (AmpC^wt^, AmpC^hyp^, TEM-1), a second complex biofilm experiment was conducted. Inocula were prepared as described above, with the additional use of a defined 19-species strain collection from a single peri-implantitis case^18^ (see below). The experiment was performed anaerobically and consequently the 96-well plates were incubated for 24 hours at 37°C in anaerobic conditions. After 24 hours the supernatant in the 96-well plates were analyzed with a Nitrocefin (NCF) assay as described above.

### Assembly of a 19-species inoculum from clinical isolates co-recovered from a peri-implantitis case

To generate a complex, yet defined inoculum for a biofilm experiment, we combined 19 strains representing 18 species co-recovered from a single case of severe peri-implantitis^18^. Specifically, this culturome included *Neisseria* sp. strain SPS_001, *Leptotrichia* sp. strain SPS_002, *Leptotrichia hofstadii* strain SPS_003, *Streptococcus anginosus* strain SPS_004, *Streptococcus* sp. strain SPS_005, *Streptococcus gordonii* strain SPS_007, *Eikenella corrodens* strain SPS_010, *Dialister pneumosintes* strain SPS_012, *Veillonella dispar* strain SPS_013, *Streptococcus cristatus* strain SPS_014, *Capnocytophaga leadbetteri* strain SPS_015, *Actinomyces* sp. strain SPS_016, *Gemella morbillorum* strain SPS_020, *Prevotella buccae* strain SPS_021, *Prevotella nigrescens* strain SPS_022, *Fusobacterium nucleatum* subsp. *vincentii* strain SPS_023, *Prevotella veroralis* strain SPS_024 and *Selenomonas* sp. strain SPS_025. Individual strains were revitalized from glycerol stocks and cultured anaerobically on MSPS_029 medium at 37°C for three days. Biomass from each strain was collected from plates and adjusted to an OD_600 nm_ of 1 in BHI supplemented with 25% glycerol. Equal volumes of each strain were mixed, aliquoted, and stored at -80°C for future use.

### DNA isolation and full-length 16S rRNA gene amplicon sequencing

The composition of biofilms was assessed using PacBio full-length 16S rRNA gene amplicon sequencing, following a previously established method^103^. In summary, DNA was extracted from the biofilms utilizing the Fast DNA Spin Kit for Soil (MP Biomedicals Germany GmbH, Eschwege, Germany). DNA concentration was then determined with the Invitrogen Qubit dsDNA BR Assay Kit and measured using the Qubit 2.0 fluorometer (Thermo Fisher Scientific, Waltham, MA, USA). Each PCR reaction was set up in a 50 µL volume, including the DNA template (standardized to 5 ng when possible), KAPA PCR mix, primers 27F (AGRGTTYGATYMTGGCTCAG) and 1492R (RGYTACCTTGTTACGACTT), along with molecular-grade water. The 16S rRNA gene target was amplified in a single PCR protocol using 25 cycles. The thermal cycling conditions began with an initial denaturation at 95°C for 3 minutes, followed by 23 to 30 cycles of denaturation at 95°C for 30 seconds, annealing at 55°C for 30 seconds, and extension at 72°C for 90 seconds. A final extension step was carried out at 72°C for 10 minutes. To evaluate PCR product integrity, 5 µL of the reaction was analyzed via agarose gel electrophoresis. On the same day, PCR products were purified using the MiniElute PCR Purification Kit 250 (Qiagen, Hilden, Germany), and DNA concentration was measured again using the Qubit dsDNA HS Assay Kit (Thermo Fisher Scientific). The purified PCR products were subsequently submitted for PacBio Sequel sequencing and analyzed using an in-house bioinformatics pipeline^103^.

### Statistical analysis

The quality-controlled output of full-length 16S rRNA gene amplicon sequencing included a datasheet detailing species or OTU counts for each biofilm sample, which were then used for multivariate statistical analyses^104, 105^. Non-metric multidimensional scaling (nMDS) was conducted based on a Bray–Curtis similarity matrix derived from biofilm composition data. Prior to analysis, the dataset was standardized by normalizing read counts to the total number of reads per sample. Taxonomic reads were then aggregated at both genus and class levels. To illustrate correlations between biofilm composition and ordination axes, vector overlays were applied. Each vector originated from the plot’s center (0, 0) and extended to coordinates (x, y), representing the Pearson correlation coefficients of the variable with ordination axes 1 and 2, respectively. The vector’s length and direction reflected the strength and orientation of the variable’s association with the axes. Permutational multivariate analysis of variance (PERMANOVA) was employed to assess the influence of inoculum and culture conditions —such as inoculum origin, presence of helper strains, supplementation of antibiotics, or phage presence—on Bray–Curtis dissimilarities in a factorial ANOVA-like design using permutation testing. Post hoc pairwise comparisons between factor levels were conducted. Species-level diversity indices were calculated using the DIVERSE function applied to rarefied abundance data.

### Electron microscopy

Virions were analyzed with electron microscopy as previously described^106^. Briefly, virions were sedimented from cooled, sterile-filtered conditioned media by ultracentrifugation at 25,000 × g for 90 min. The supernatant was discarded and replaced with 0.1 M ammonium acetate (pH 7), followed by another centrifugation. This process was repeated twice. The concentrated phages were then adsorbed onto a carbon film for 30 – 120 seconds, rinsed with molecular-grade water, and negatively stained with 3% (w/v) phosphotungstic acid (pH 7.0, Sigma-Aldrich, Germany). The samples were mounted on a 400-mesh copper specimen grid (Plano, Germany) and examined using a Morgagni transmission electron microscope (FEI, U.S.) at an acceleration voltage of 80 kV.

## Data availability

The 16S rRNA gene amplicon sequencing data have been deposited in NCBI under BioProject ID: PRJEB1270987 and will be made available upon article release. All other data are currently accessible upon reasonable request and will likewise be shared at the time of publication.

## Acknowledgments

This study was funded by the Deutsche Forschungsgemeinschaft (DFG, German Research Foundation) – SFB/TRR-298-SIIRI – Project-ID 426335750 (SPS and MSti). Additional support for this study was provided by the DFG under Germany’s Excellence Strategy (EXC 2155, Project Number 390874280, awarded to MSti). MSte would like to thank Ministry of Science and Culture of Lower Saxony (Niedersächsisches Ministerium für Wissenschaft und Kultur) for funding through BacData, ZN3428 awarded to MSti. IY is funded by the “Federal and State Program Promoting Female Professors”, Grant No. 01FP19068J. We also thank Diana Strauch and Rainer Schreeb for their valuable technical assistance, and Wiebke Behrens for support in performing Pacbio sequencing.

