## Supplementary material for "Invasive species drive polymicrobial resistance to amoxicillin in oral biofilms through β-lactamase release": SI

Short title: Cross-protection in biofilms

Amy L. Seidel[1,2], Matthias Steglich[1,2], Taoran Qu[1,2], Amruta A. Joshi[1,2], Malina Schneider[1,2], Andrea Spaic[1,2], Ines Yang[1,2], Jasmin Grischke[1,2], Evgenii Rubalskii[2,3], Lothar Koch[2,4], Boris Chichkov[2,4], Jan Hegermann[5], Piotr Majewski[6], Elżbieta Tryniszewska[6], Jørgen Slots[7], Szymon P. Szafrański[1,2,8,*,$] and Meike Stiesch[1,2,8,*,$]

[1]Department of Prosthetic Dentistry and Biomedical Materials Science, Hannover Medical School, Hannover, Germany

[2]Lower Saxony Centre for Biomedical Engineering, Implant Research and Development (NIFE), Hannover, Germany

[3]Department of Cardiothoracic, Transplantation and Vascular Surgery, Hannover Medical School, Hannover, Germany.

[4]Institute of Quantum Optics, Leibniz Universität Hannover, Hannover, Germany

[5]Research Core Unit Electron Microscopy, Institute of Functional and Applied Anatomy, Hannover Medical School, Hannover, Germany

[6]Department of Microbiological Diagnostics and Infectious Immunology, Medical University of Bialystok, Poland

[7]Division of Periodontology, Diagnostic Sciences and Dental Hygiene, Ostrow School of Dentistry of USC, University of Southern California, Los Angeles, California, USA

[8]Cluster of Excellence RESIST (EXC 2155), Hannover Medical School, Hannover, Germany

[*]Szymon P. Szafrański and Meike Stiesch contributed equally

[$]correspondence to: Meike Stiesch and Szymon P. Szafrański, Department of Prosthetic Dentistry and Biomedical Materials Science, Hannover Medical School, Carl-Neuberg-Str. 1, 30625 Hannover, Germany

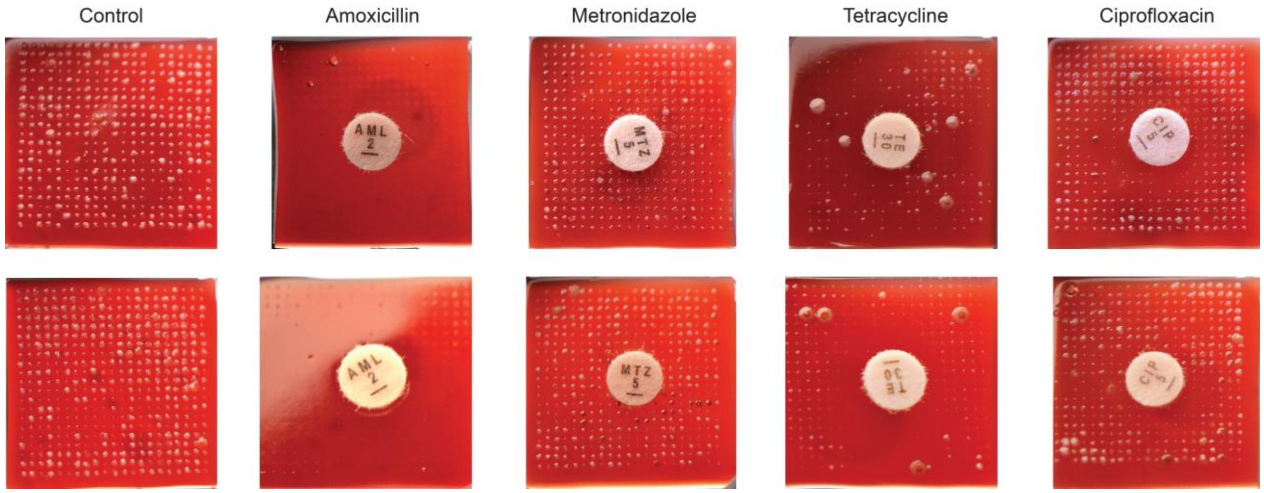

**Fig. S1.** Example bioprinted biofilm arrays for antimicrobial susceptibility testing of oral biofilms. Control biofilms and colony biofilms around disks with four different antibiotics are shown for two implants.

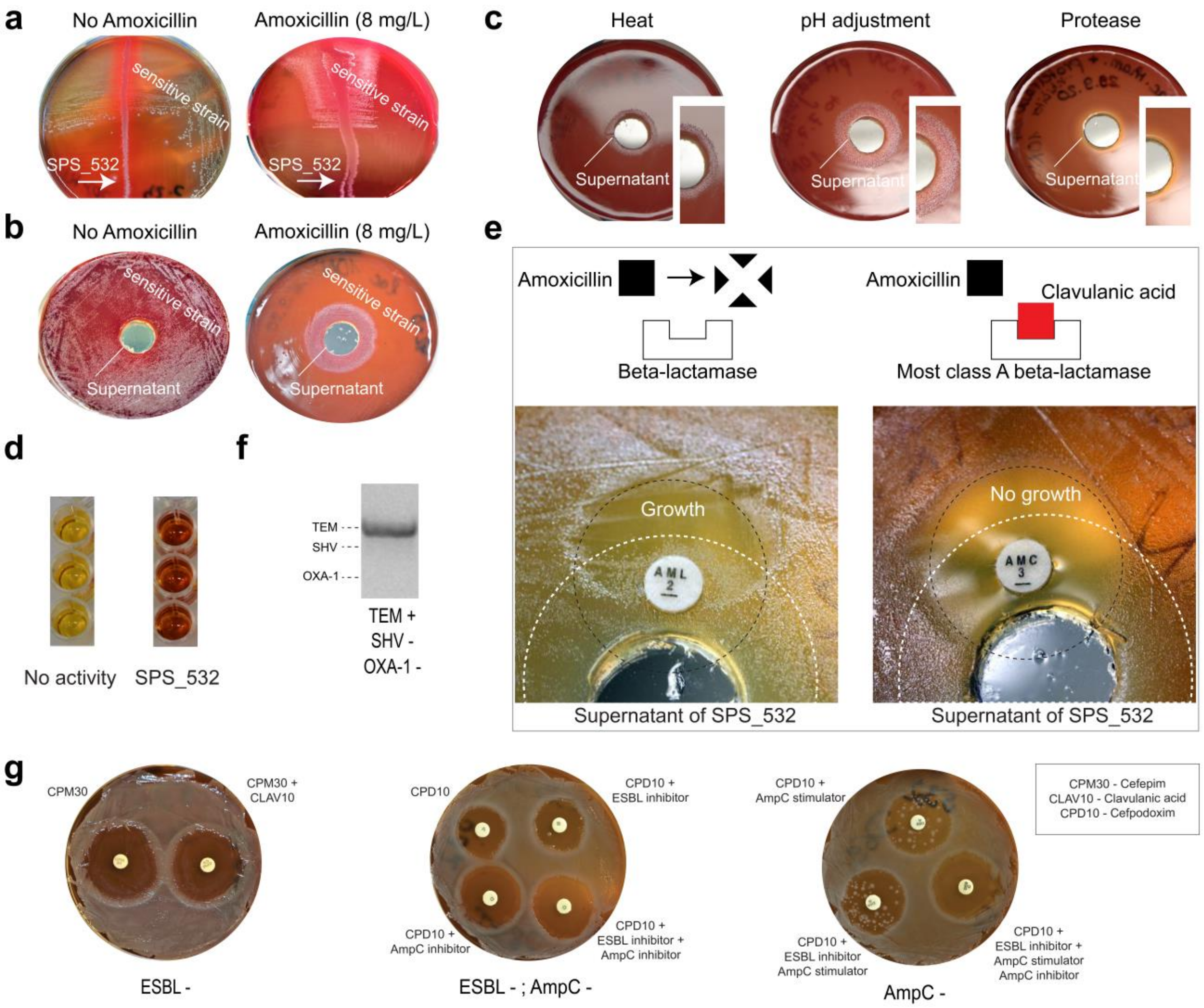

**Fig. S2**. Mechanism of cross-protection in a peri-implantitis case.

**a** The cross-protection was reproduced using isolates from bioprints arrays with satellite growth. **b** The cross-protection was conferred by spent medium from a helper strain culture. **c** Effect of different treatments on spent medium activity. **d** Nitrocefin assay for detection of β-lactamase activity. **e** Effect of clavulanic acid on cross-protection activity. **f** PCR-based detection of *bla*$_{TEM}$, *bla*$_{SHV}$ and *bla*$_{OXA}$ genes. **g** Three disks assays for detection of AmpC hyper-production or ESBLs or both in strains SPS_532.

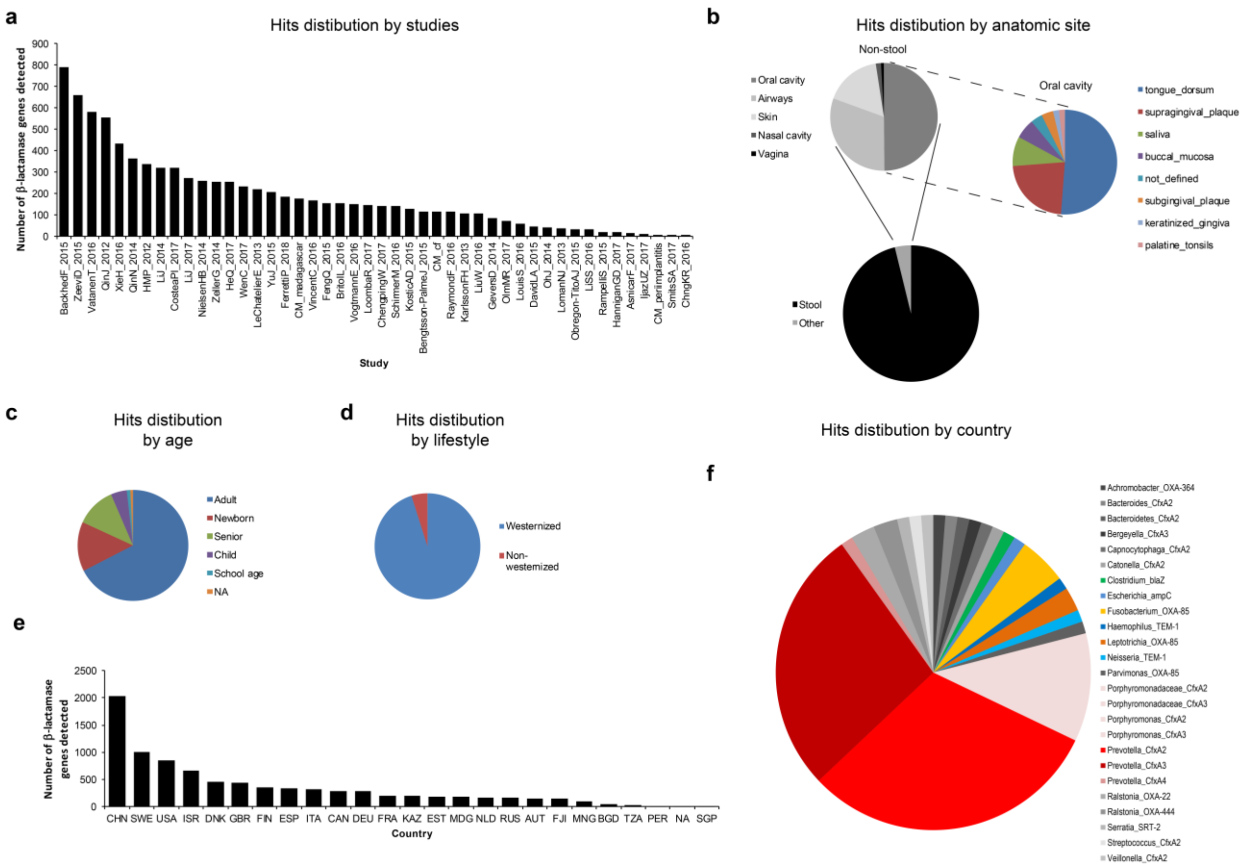

**Fig. S3.** β-lactamase genes in human-associated metagenomes

**a** Incorporated original studies. **b** Distribution of anatomical sites. **c** Age distribution. **d** Lifestyle distribution. **e** Country distribution. **f** Major oral producers and their enzymes.

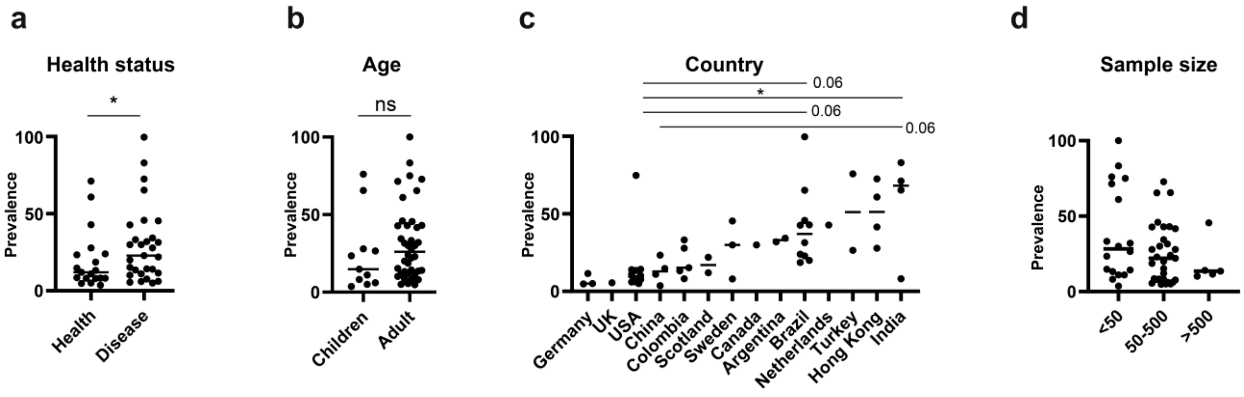

**Fig. S4.** Prevalence of enteric Gram-negative rods across populations' characteristics.

**a** Health status. **b** Age. **c** Geographical location. **d** Size of the study.

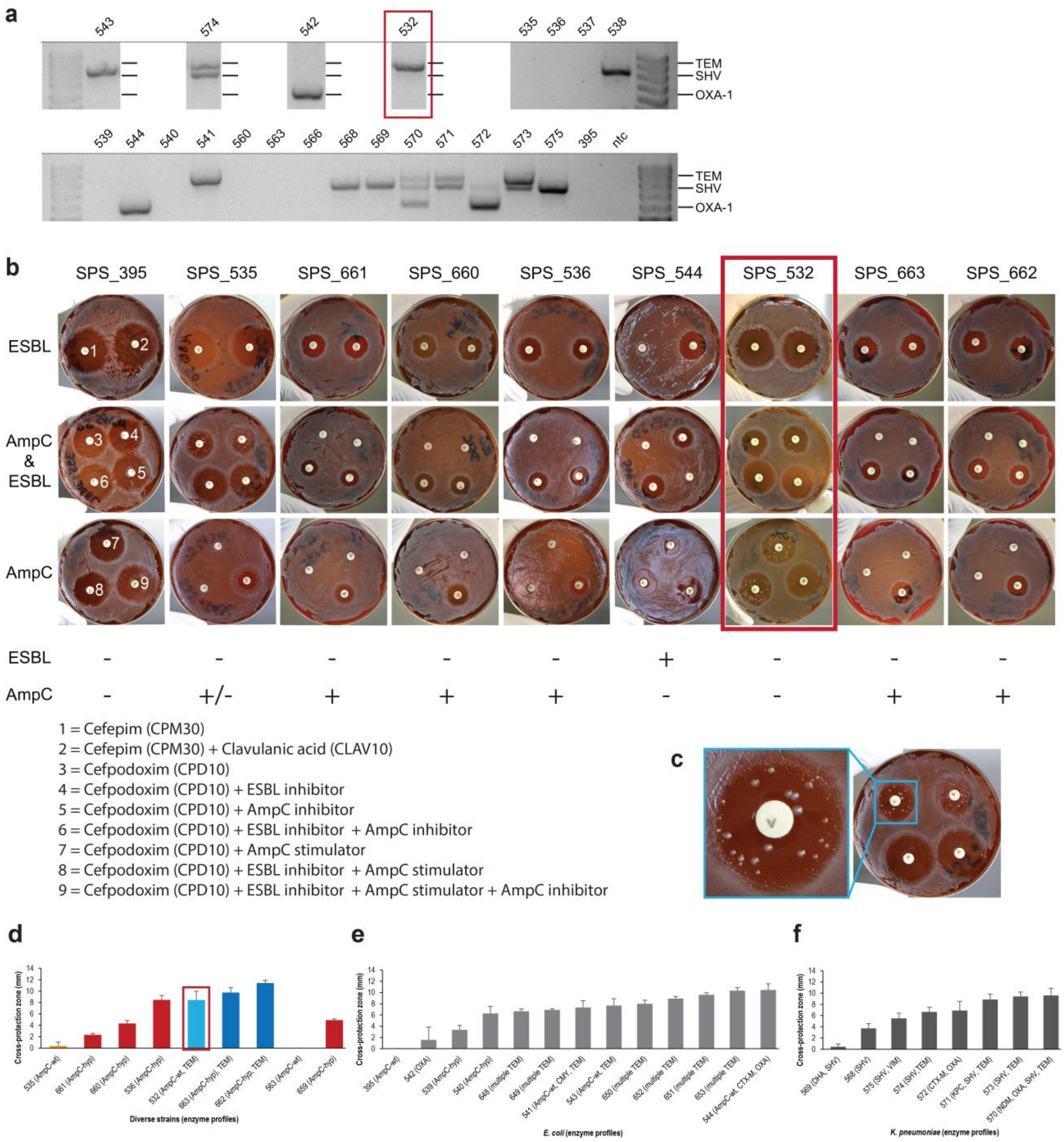

**Fig. S5.** Strains conferring cross-protection from Amoxicillin.

**a** PCR-based characterization of β-lactamase genes in strains. An example detection of *bla*$_{TEM}$, *bla*$_{SHV}$ and *bla*$_{OXA}$ genes. **b** Three disk assays for detection of AmpC hyper-production or ESBLs or both. **c** Colonies of AmpC hyper-producer mutants deriving from *Enterobacter* sp. strain SPS_532 around Cefpodoxime disks. **d** Cross-protecting mutants characterized by AmpC hyper-production. Examples for *Escherichia coli* (Ec), *Enterobacter* sp. (Es) and *Klebsiella aerogenes* (Ka) are shown. **e** Cross-protection activity across selected *E. coli* strains. **f** Cross-protection activity across selected *Klebsiella pneumoniae* strains

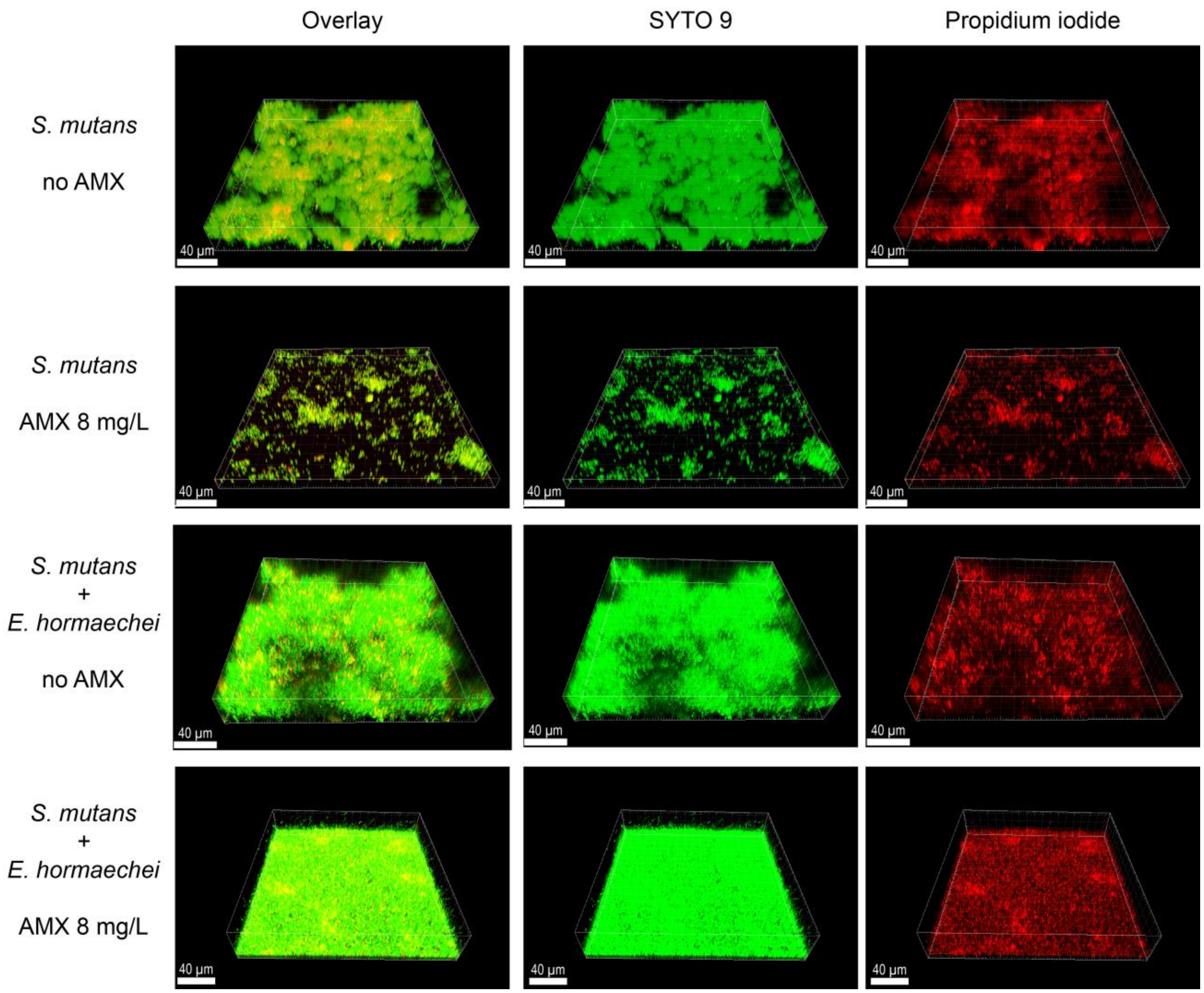

**Fig. S6.** Cross-protection in two-species model.

Effect of Amoxicillin and presence of *Enterobacter* sp. strain SPS_532 on the biofilm formation of *Streptococcus mutans*. CLSM micrographs are shown for life (SYTO) and dead (Propiodium iodide) cells, as well as as an overlay.

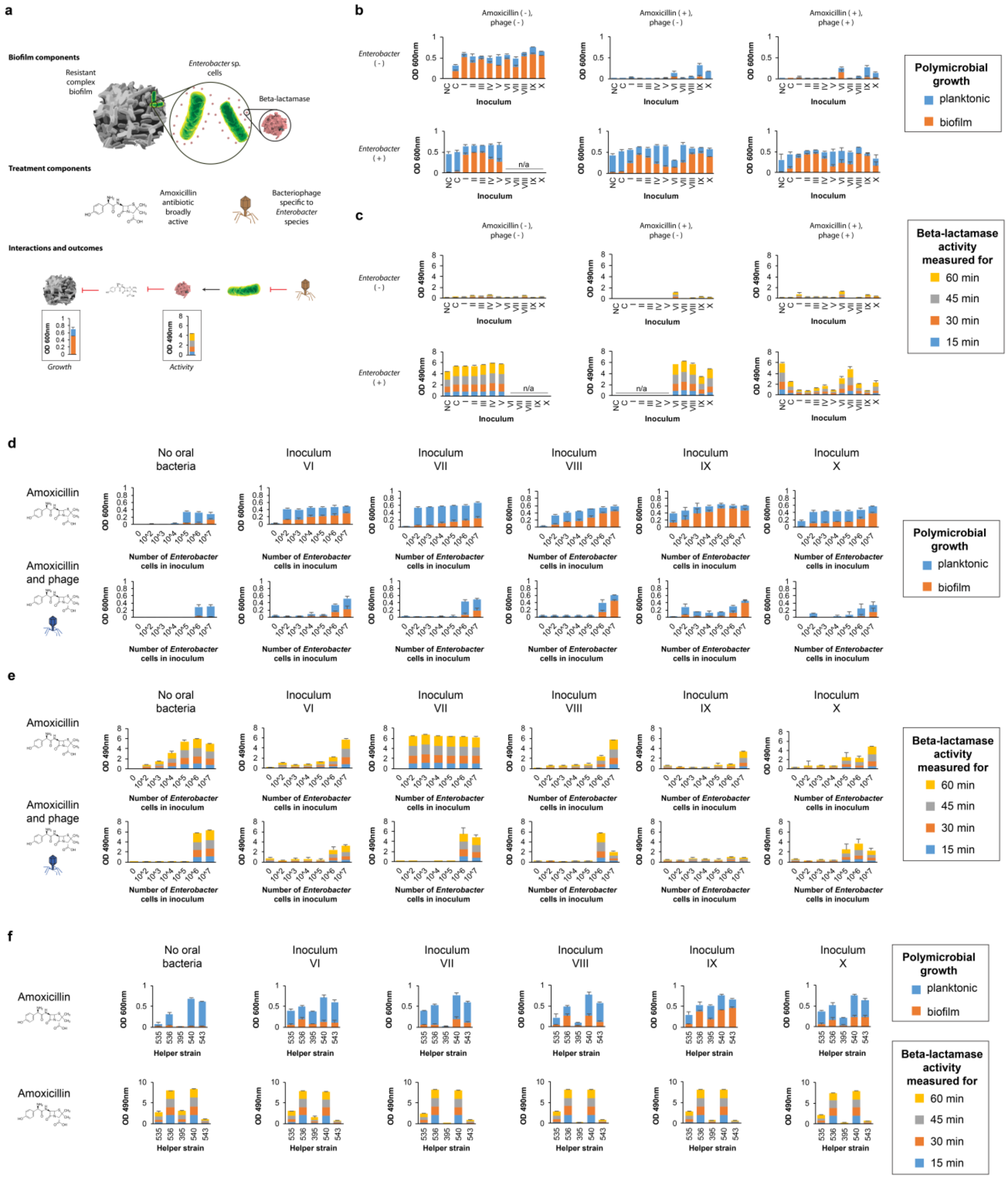

**Fig. S7.** Cross-protection in complex biofilms and its phage-based control across complex biofilm inocula.

**a** Framework of the complex biofilm model. **b** Relationship between complex biofilm growth and presence of helper in two treatment groups. **c** Relationship between β-lactamase activity and presence of helper in two treatment groups. **d** Relationship between complex biofilm growth and size of helper's inoculum in two treatment groups across inocula. **e** Relationship between β-lactamase activity and size of helper's inoculum in two treatment groups across inocula. **d** Effect of AmpC hyperproduction and TEM production on complex biofilm growth (top) and β-lactamase activity (bottom) across inocula.

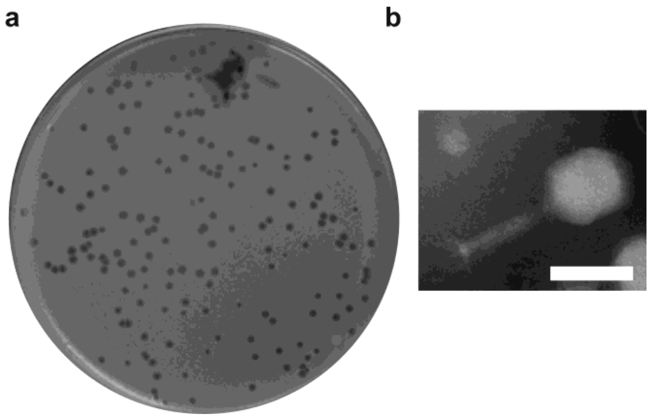

**Fig. S8.** Characteristics of *Enterobacter* phage used in complex biofilm model

**a** Plaque morphology. Plate has diameter of 100 mm. **b** Virion morphology (Courtesy of Dr. Jan Hegermann, Hannover Medical School). Bar represents 100 nm.

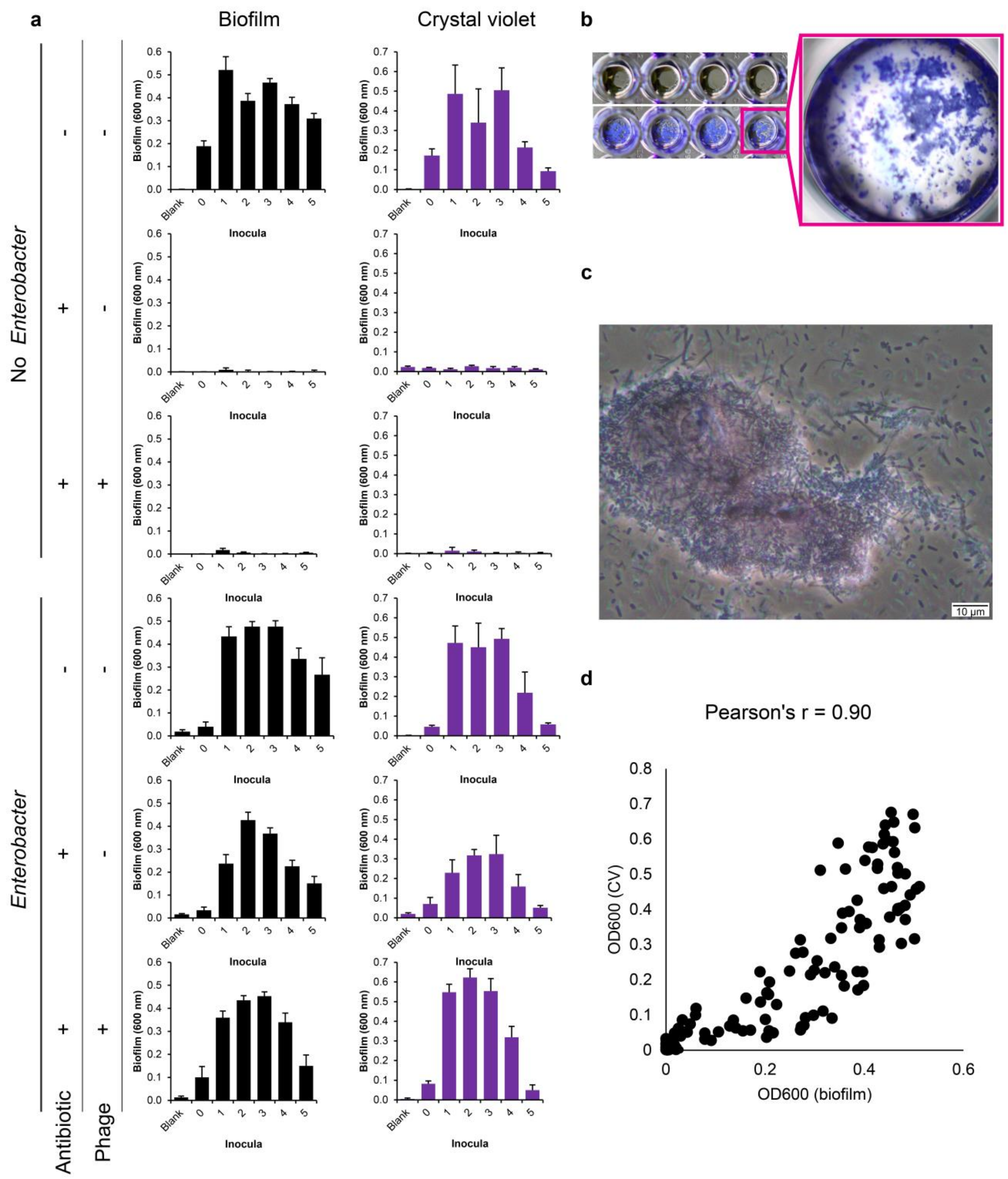

**Fig. S9.** Relationship between direct $OD_{600nm}$ and crystal violet $OD_{600nm}$ measurements for complex biofilms. **a** Direct $OD_{600nm}$ and crystal violet $OD_{600nm}$ measurements for six experimental groups and six inocula. **b** Photograph of a representative crystal-violet-stained biofilm in the microtiter plate wells. **c** Micrograph of crystal-violet-stained biofilm fragment. **d** Correlation between direct $OD_{600nm}$ and crystal violet $OD_{600nm}$ measurements.

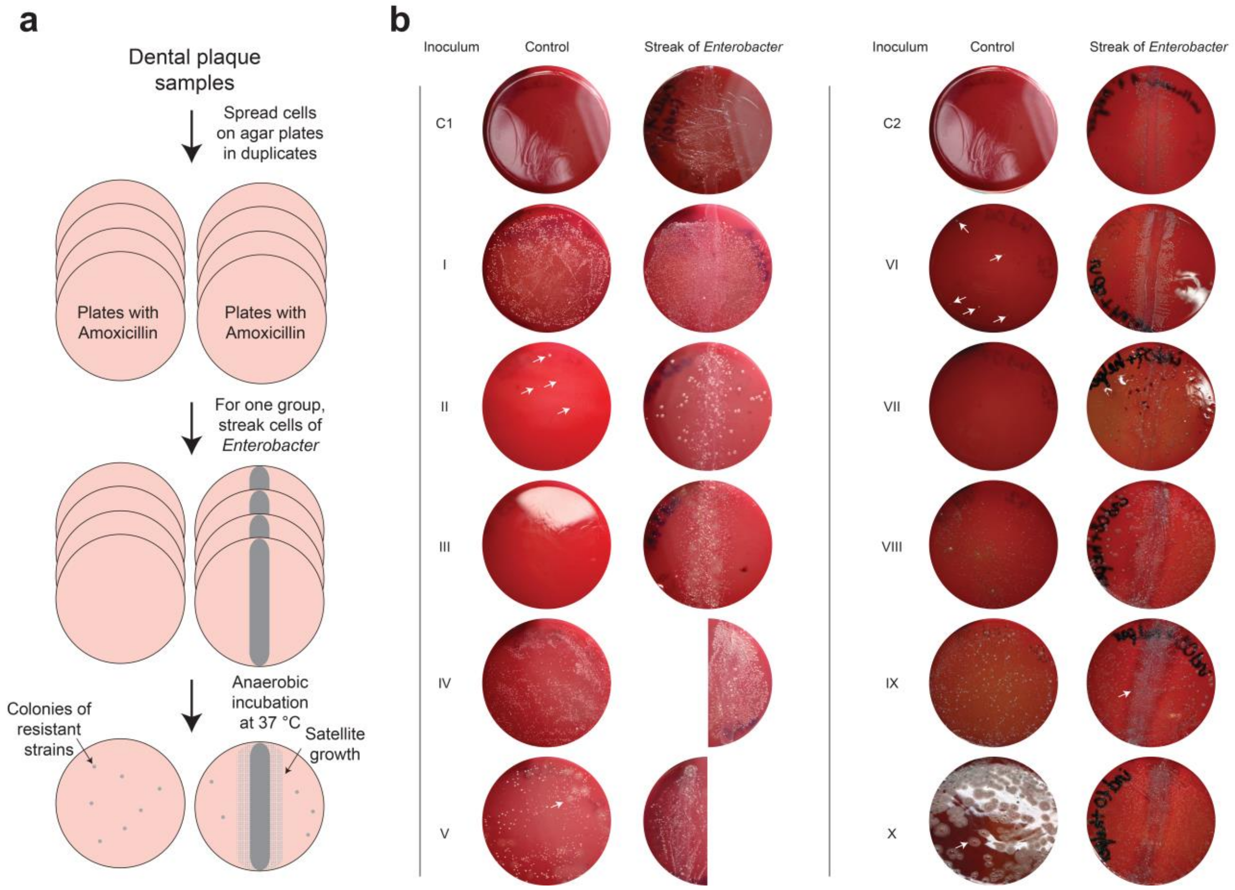

**Fig. S10.** Diffusion agar plate assay for cross-protection between a helper strain and complex biofilms.

**a** A workflow. **b** Results for a complex 19-species biofilm (C1 and C2 represent two biological replicas) and ten complex clinical inocula.

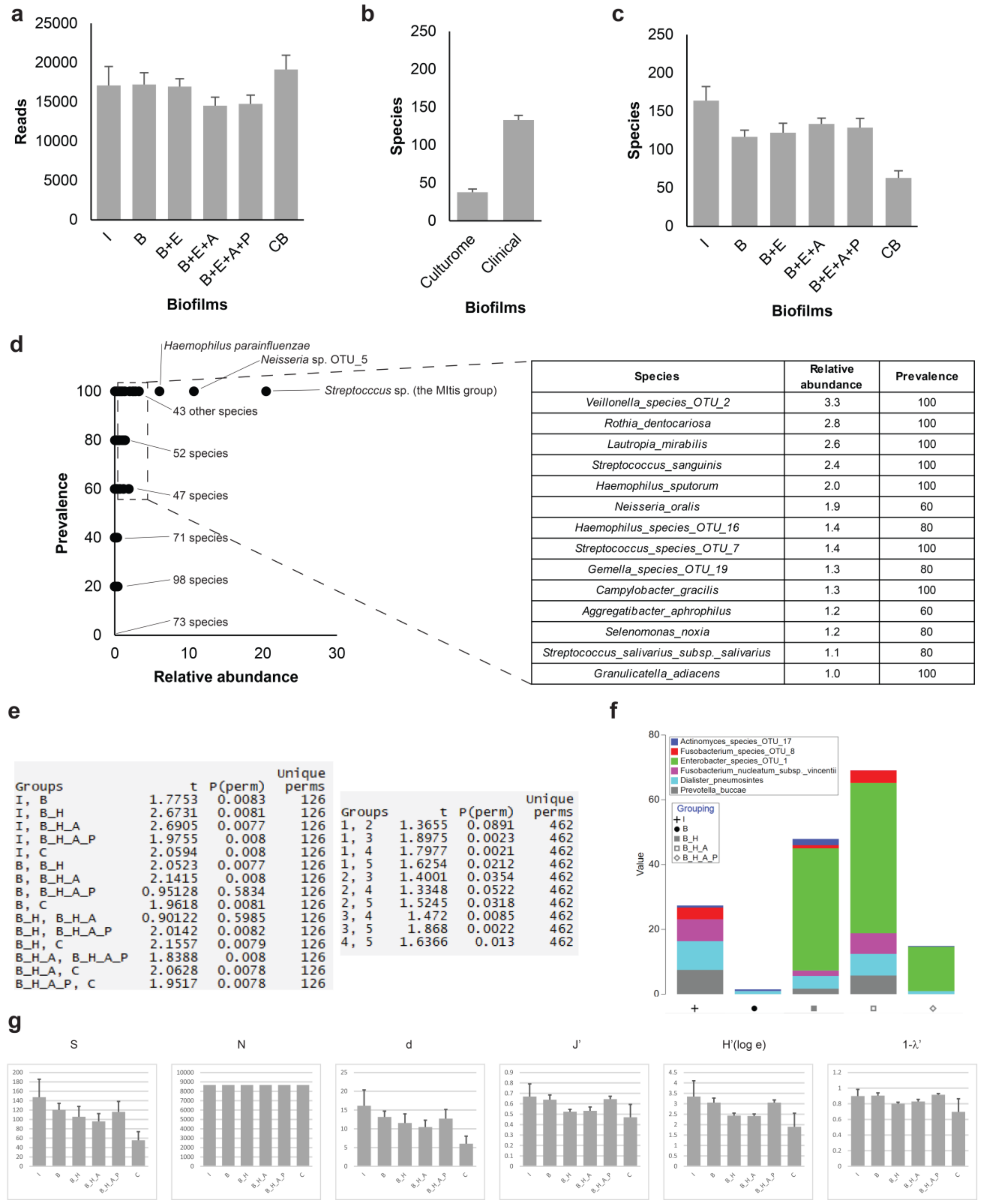

**Fig. S11.** Effect of cross-protection and phage treatment on composition of complex biofilms; additiona l information.

**a** Sequencing depth of Pacbio SMRT sequencing of 16S rRNA gene amplicons showed for experimental groups. The groups consist of inocula (I), *in vitro* biofilms (B), biofilms supplemented with *Enterobacter* helper (B+E), and additionally with Amoxicillin (B+E+A), and additionally with *Enterobacter* phage (B+E+A+P) as well as satellite colony biofilms formed on medium with Amoxicillin around *Enterobacter* streak in diffusion assay (CB). **b** Species richness achieved using defined complex inculum compared to complex clinical inoculum. **c** Species richness showed for experimental groups. **d** Composition of complex inocula. **e** Differences in compostion between experimental groups and inocula, PERMANOVA results. P(perm), P-values; Unique perms, numbers of unique permutations. **f** Relative abundance of selected species across experimental groups in biofilms inoculated with a defined strain collection. **g** Diversity indices across experimental groups.
